# Deep generative modeling of temperature-dependent structural ensembles of proteins

**DOI:** 10.1101/2025.03.09.642148

**Authors:** Giacomo Janson, Alexander Jussupow, Michael Feig

## Abstract

Deep learning has revolutionized protein structure prediction, but capturing conformational ensembles and structural variability remains an open challenge. While molecular dynamics (MD) is the foundation method for simulating biomolecular dynamics, it is computationally expensive. Recently, deep learning models trained on MD have made progress in generating structural ensembles at reduced cost. However, they remain limited in modeling atomistic details and, crucially, incorporating the effect of environmental factors. Here, we present aSAM (atomistic structural autoencoder model), a latent diffusion model trained on MD to generate heavy atom protein ensembles. Unlike most methods, aSAM models atoms in a latent space, greatly facilitating accurate sampling of side chain and backbone torsion angle distributions. Additionally, we extended aSAM into the first reported transferable generator conditioned on temperature, named aSAMt. Trained on the large and open mdCATH dataset, aSAMt captures temperature-dependent ensemble properties and demonstrates generalization beyond training temperatures. By comparing aSAMt ensembles to long MD simulations of fast folding proteins, we find that high-temperature training enhances the ability of deep generators to explore energy landscapes. Finally, we also show that our MD-based aSAMt can already capture experimentally observed thermal behavior of proteins. Our work is a step towards generalizable ensemble generation to complement physics-based approaches.

## INTRODUCTION

Machine learning (ML) has made immense strides forward in modeling 3D structures of proteins, with predictions that closely approximate ensemble-averaged experimental structures. Thanks to the pioneering work of AlphaFold2^1^ (AF2), structure prediction is now becoming a reliable technique with relevance in biomedical sciences^2^. New generations of models, like AlphaFold3 (AF3), have further pushed the boundaries, generalizing biomolecular modeling to protein complexes with diverse molecular species^3,4^.

Like all biomolecules, proteins are dynamic and exist in populations of conformations, forming structural ensembles^5^. Characterizing these ensembles is highly relevant to understand and design^6^ biological activity, yet it remains challenging both experimentally and computationally. As popular structure predictors struggle at capturing protein dynamics^7^, novel deep learning methods are being proposed to tackle the challenge^8^. Current approaches differ in their objectives and methodology. Some methods attempt to capture conformational variability as observed in the Protein Data Bank^9^ (PDB). Their goal is to sample alternative experimental-like states of proteins^10^ (e.g.: apo-holo, open-closed states). Another class of methods, including our work, follows a different approach by training on simulation data, such as molecular dynamics (MD). MD is an invaluable methodology, thanks to its ability to capture physical behavior of biomolecules^11,12^. Unfortunately, the large and rugged energy landscapes of biomolecules allow MD to be effective only at enormous computational cost^13–15^ or requiring specialized hardware^16^.

Given their speed, ML methods trained on MD are a promising strategy to provide a fast approach to generating conformational ensembles^17^. Research in this direction has started to flourish in the post-AF2 era, thanks to three key factors: (i) the neural network components from AF2 have proven highly effective at modeling protein structure and are often directly reused in MD ensemble generators^18^; (ii) advancements in deep generative modeling, especially diffusion or flow matching models^19,20^, have been successfully repurposed for this task; (iii) the availability of large MD datasets^21,22^ has made it possible to train transferable models that can be applied to proteins outside training data. One model pioneering this paradigm is AlphaFlow^18^, an AF2 version re-wired as a generative model. The method was trained on the ATLAS dataset^22^ of simulations of protein chains from the PDB. It was shown to accurately reproduce some MD properties, such as residue fluctuations, but appeared to fail at capturing more complex multi-state ensembles in that database^23^. Similar methods have also been trained on ATLAS^24^ or comparable datasets^25^. Recently, the BioEmu model^26^ demonstrated for the first time that an MD-based generative model can consistently capture alternative states of proteins outside its training data. BioEmu is a diffusion model harnessing neural networks components from AF2 and was trained on a massive MD dataset that is not fully available. BioEmu represents a milestone in structural ensemble modeling via ML but still presents limits. For example, it only directly generates backbone atoms of proteins and to model side chains it needs a post-processing step with a third-party neural network followed by energy minimization. Additionally, it models structural ensembles only at 300 K, the predominant simulation temperature in its training data.

Given the progress in modeling proteins in fixed conditions, one of the next challenges for ML generators is now to achieve physical transferability to different environmental conditions. For instance, temperature is a fundamental thermodynamic parameter that influences conformational ensembles according to the Boltzmann distribution, primarily by shifting the balance between enthalpy and entropy. The effect of temperature is most clearly seen as a determinant of protein folding^27^, directly impacting biological function^28^. Also, the thermal behavior of proteins is relevant in biotechnology and evolutionary biology^29,30^.

Here, we present a deep generative model that directly produces heavy atom ensembles of proteins under varying environmental conditions. The method is an evolution of the Structural Autoencoder Model (SAM), a latent diffusion model^31^ that we originally applied to Cα ensembles of intrinsically disordered peptides^32^. We first compare the new aSAM (atomistic SAM) with the state-of-the-art AlphaFlow on the ATLAS dataset. Our model achieves similar performance to AlphaFlow without relying on a pre-trained AF2 version and with higher computational efficiency when generating ensembles. Moreover, we show that modeling atoms in a latent space with aSAM is an effective strategy for learning physically realistic distributions of backbone and side chain torsion angles, a challenge that AlphaFlow and most existing MD ensemble generators struggle with.

Importantly, we also introduce aSAMt (aSAM temperature-conditioned), a proof-of-principle ML generator that produces protein conformational ensembles conditioned by a physical variable, in this case, temperature. For training, we leveraged the large and openly available mdCATH dataset^21^, which contains MD simulations for thousands of globular protein domains at different temperatures, from 320 to 450 K. We show that aSAMt can recapitulate the temperature behavior of a variety of ensemble properties and that it is able to generalize in a meaningful way not just to unseen sequences and structures but also to temperatures outside its training data. Furthermore, by comparing its ensembles with long MD simulations of fast folding proteins^12^, we show that learning from high temperatures greatly enhances the capability of ML ensemble generators to explore conformational landscapes. Thanks to the multi-temperature mdCATH training data, our model obtains comparable coverage of the energy landscapes of these fast-folding proteins with respect to BioEmu, despite not being directly trained on their extensive simulations. Lastly, we show how, by using information learned solely from MD, the model can capture experimentally observed temperature behavior, establishing how learning from simulation data can be a fruitful pre-training strategy to model experimental observations. Taken together, aSAM represents a significant contribution in the field of modeling protein structural ensembles with deep learning.

## RESULTS

### aSAM, a latent diffusion model for atomistic ensemble generation

aSAM includes two components: an autoencoder (AE) is initially trained to represent heavy atom coordinates of proteins as SE(3)-invariant encodings (**Fig. 1a**). Then, a diffusion model is applied to learn the probability distribution of such encodings (**Fig. 1b**). The latter model is conditioned on an initial 3D structure representing the conformation from which an MD simulation would start. In the aSAM version trained on mdCATH (aSAMt), it is also conditioned on temperature. The generation of an ensemble is performed by sampling encodings via the diffusion model and mapping them to 3D structures with the decoder (**Fig. 1c**). In the decoder, we leverage the Structure Module from AF2 for heavy atom generation. Because this module treats most bond lengths and angles as rigid, it decreases the number of degrees of freedom to learn.

**Fig. 1.**
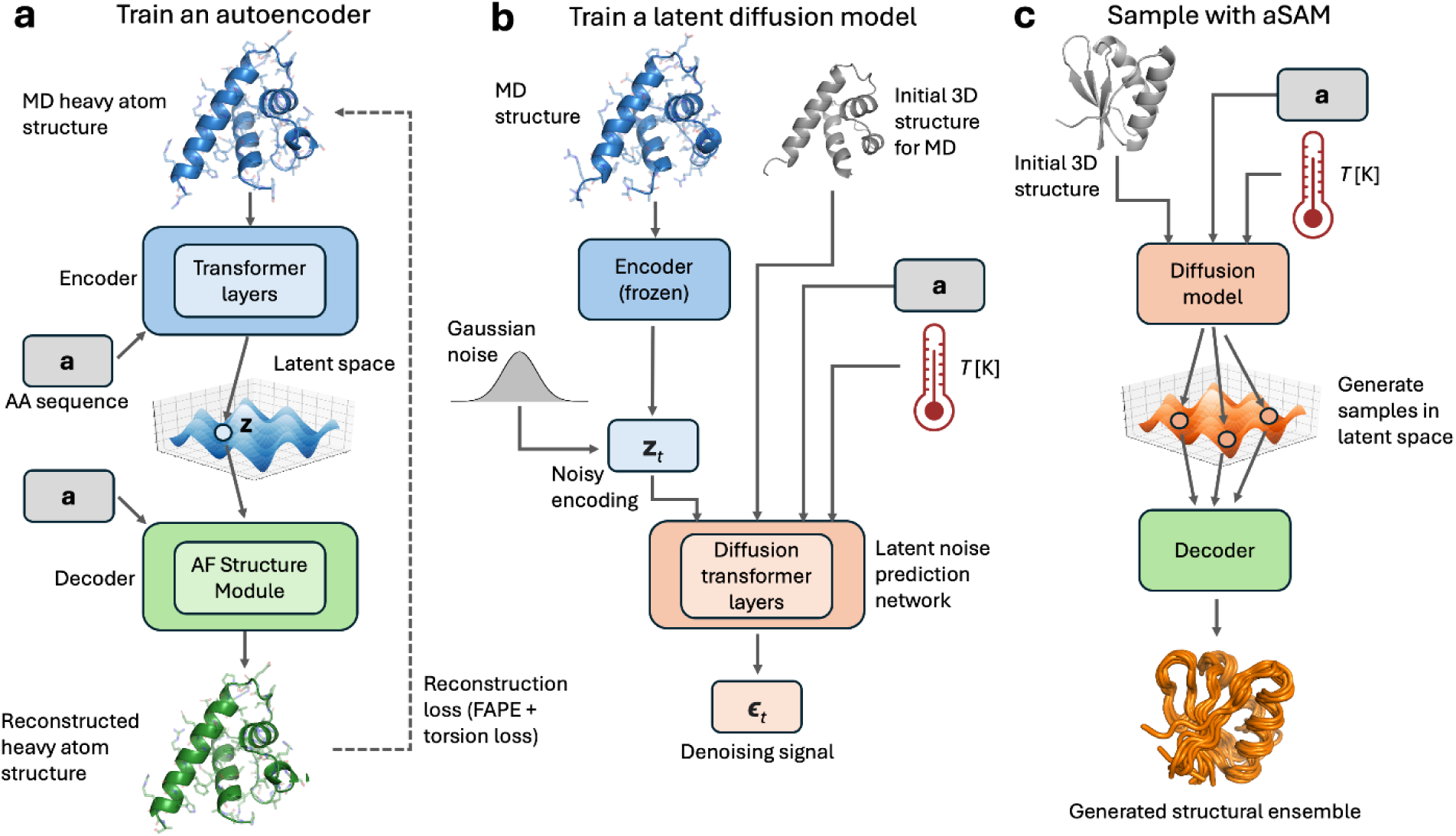
Overview of aSAM. **a** In the first stage of aSAM training, an AE learns to encode heavy atom protein structures into a SE(3)-invariant latent space. **b** In the second stage, a denoising probabilistic diffusion model learns the distribution of MD structures in the latent space. The encoder is not trainable anymore at this stage. The diffusion model is conditioned on an initial MD structure, its amino acid sequence and (optionally) a temperature scalar value. **c** Sampling with aSAM at inference time. The diffusion model samples from the latent space and the decoder maps the generated elements to 3D structures to obtain 3D structural ensembles.

A key requirement for latent diffusion models is a good reconstruction ability of the decoder^32,33^. In this work, we used two MD datasets, ATLAS and mdCATH (see below), and we trained a separate AE on each. To assess the decoders, we evaluated them on MD snapshots of test set proteins from both datasets. The decoders typically reconstruct encoded conformations with a heavy atom root mean square deviation (RMSD) of 0.3-0.4 Å (**Supplementary Table 1**), corresponding to highly accurate reconstructions.

While the aSAM decoders work well for MD snapshots, the diffusion model may generate encodings that do not exactly map onto stereochemically correct configurations. As a result, such encoding may reconstruct globally correct 3D structures but with atom clashes, especially for side chains. This is likely caused by the limited capacity of our diffusion model to learn high-resolution atomistic details in the latent space. To avoid clashes, deep learning methods may apply physics-based terms in loss functions, like in AF2^1^, or add specialized layers, like atom-transformers from AF3^3^. In this work, we address the problem by briefly relaxing the 3D structures from aSAM with a computationally efficient custom energy minimization protocol (**Methods**). We found it effective at removing heavy clashes without perturbing backbone atoms significantly, as measured by average backbone RMSDs ranging from 0.15 to 0.60 Å after minimization, depending on the dataset (**Supplementary Fig. 1**). Unless otherwise stated, all results in this work originate from aSAM sampling followed by energy minimization.

### Benchmarking aSAM on ATLAS

As a first validation of aSAM, we compared it with AlphaFlow, a state-of-the-art ensemble generator. Since AlphaFlow was trained on the ATLAS dataset, we trained an aSAM version on the same data. As ATLAS simulations are all performed at 300 K, this model is not temperature-conditioned and we named it aSAMc (aSAM constant-temperature). We focused on the template-based version of AlphaFlow which, like SAM, is conditioned on an input initial structure. The same input was provided to the two models.

Both methods provided good approximations of MD ensembles for a fraction of test set proteins, typically the ones with rigid structures and a few short flexible elements, such as 6h49_A and 7e2s_A (**Fig. 2a and b**, and **Supplementary Fig. 2a** and b). In these cases, the models capture local dynamics, as observed by Cα root mean square fluctuation (RMSF) profiles. They also cover most of the diversity observed in reference MD simulations, as seen by the distribution of Cα RMSD to the initial structure of MD (initRMSD), and principal component analysis (PCA) of the ensembles. However, for proteins with more complex multi-state ensembles, such as 6q9c_A (**Fig. 2e and f**), or for proteins with long flexible elements that broadly sample the conformational landscape, like 7c45_A (**Supplementary Fig. 2e** and f), both models fail to capture states distant from the initial structure.

**Fig. 2.**
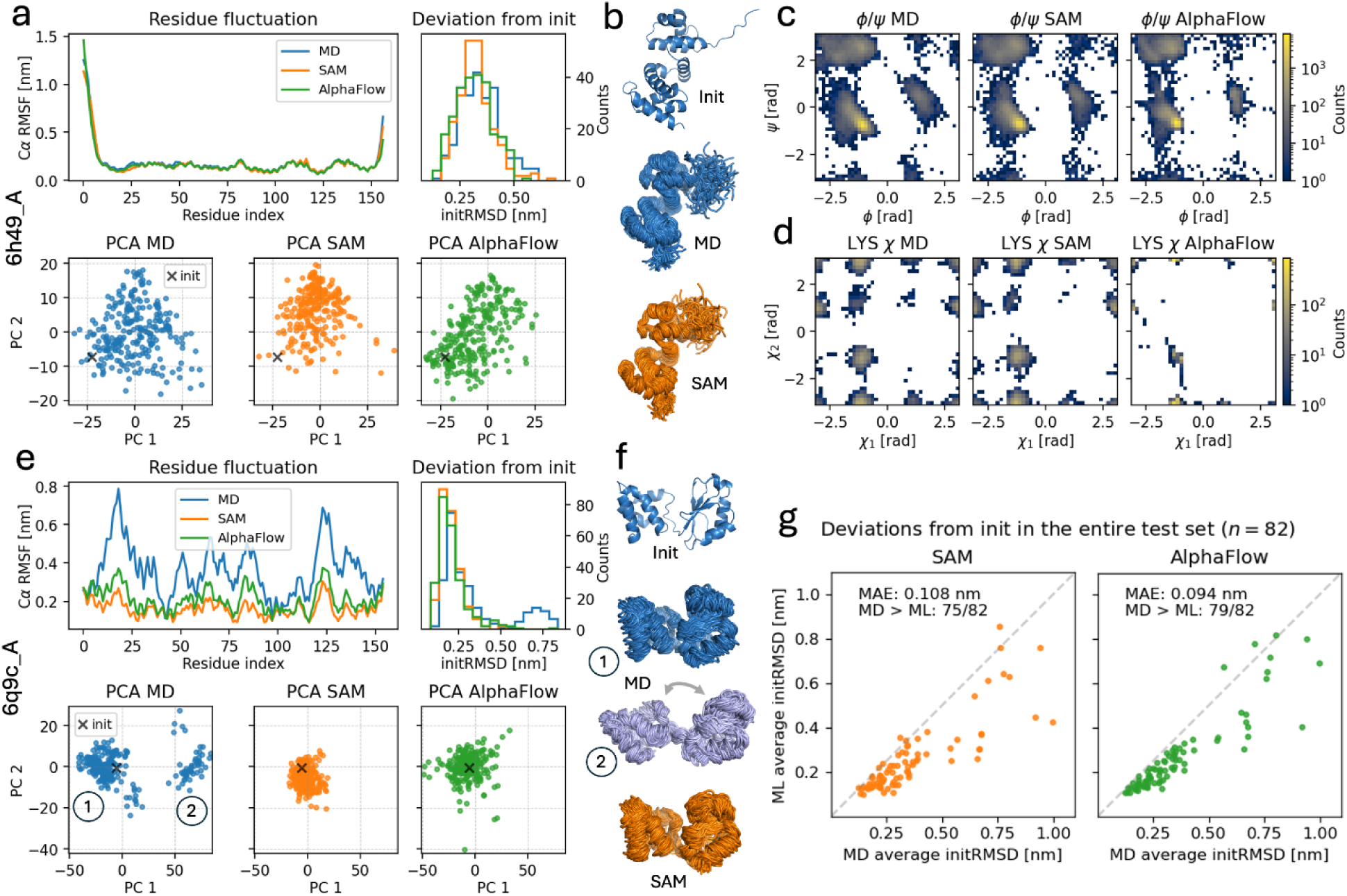
aSAMc and AlphaFlow ensembles for ATLAS test set proteins. **a** Properties of MD (blue), aSAMc (orange) and AlphaFlow (green) ensembles of test chain 6h49_A, including Cα RMSF profiles and initRMSD histograms (top), and PCA projections (bottom). The black × marks the initial MD structure input to ML models. **b** Rendering of the initial MD structure and ensembles of 6h49_A. **c** *ϕ* and *ψ* torsion angle histograms from all residues in the 6h49_A ensembles. **d** *χ*_1_ and *χ*_2_ torsion angle histograms for all lysine side chains in MD and ML ensembles of 6h49_A (lysine is chosen as a representative amino acid). **e** Analysis of the test chain 6q9c_A. The MD ensemble has a closed (1) and open (2) states. They are visible in PCA and in the initRSDM histogram as two peaks. The ML models do not sample state 2, which is geometrically distant from their input structure. **f** Initial MD structure and ensembles of 6q9c_A. aSAMc only samples the closed state. **g** Average initRMSD of MD and ML ensembles across all test set proteins (*n* = 82). The plots report the MAE of ML vs MD values and the number of proteins where ML underestimates the MD value. **a** to **g**: all data from ensembles of 250 snapshots. The 3D ensembles in **b** and **f** are subsets of 100 snapshots.

We then quantitatively evaluated the two models on the whole test set to identify systematic differences between them. We first compared their abilities to replicate local flexibility, based on the Pearson correlation coefficient (PCC) between the Cα RMSF values of MD and ML ensembles. AlphaFlow outperforms aSAMc in terms of average PCCs, with values of 0.904 and 0.886 respectively, with a statistically significant difference according to the Wilcoxon signed-rank test (**Table 1**).

**Table 1:** Evaluating ensemble generators in the reproduction of ATLAS MD ensembles.

| Strategy | PCC Cα<br>RMSF (↑) | WASCO-global (↓) | WASCO-local (↓) | chiJSD (↓) | Heavy<br>clashes (↓) | Peptide bond length<br>violations (↓) |
| --- | --- | --- | --- | --- | --- | --- |
| aSAMc<sup>a</sup> | 0.886 ± 0.011 | 158.2 ± 16.8 | 1.15 ± 0.05 | 0.07 ± 0.002 | 0.23 ± 0.04 | 0.52 ± 0.09 |
| aSAMc no-min<sup>b</sup> | 0.886 ± 0.010 | 155.5 ± 16.6 | 1.14 ± 0.05 | 0.07 ± 0.002 | 10.20 ± 1.05 | 29.04 ± 2.51 |
| aSAMc $s=50$<sup>c</sup> | 0.887 ± 0.011 | 159.1 ± 16.9 | 1.15 ± 0.05 | 0.07 ± 0.002 | 0.25 ± 0.05 | 0.54 ± 0.09 |
| aSAMt<sup>f</sup> | 0.852 ± 0.013 | 203.2 ± 25.9 | 1.47 ± 0.06 | 0.12 ± 0.002 | 0.55 ± 0.03 | 0.25 ± 0.04 |
| AlphaFlow<sup>‡</sup> | 0.904* ± 0.009 | 149.5* ± 14.9 | 1.28* ± 0.06 | 0.16* ± 0.002 | 1.14* ± 0.11 | 67.32* ± 5.82 |
| ESMFlow<sup>‡</sup> | 0.887<sup>†</sup> ± 0.011 | 157.5<sup>†</sup> ± 16.3 | 1.28* ± 0.06 | 0.16* ± 0.002 | 1.15* ± 0.12 | 58.49* ± 4.90 |
| COCOMO<sup>d</sup> | 0.798 ± 0.016 | 228.2 ± 22.8 | 2.37 ± 0.09 | 0.24 ± 0.002 | 22.12 ± 1.98 | 129.68 ± 10.33 |
| MD<sup>e</sup> | 0.998 ± 0.0002 | 45.1 ± 4.2 | 0.34 ± 0.01 | 0.01 ± 0.0001 | 0 | 0.74 ± 0.06 |

For each score, the table shows its average value and standard error across all proteins in the ATLAS test set ( $n = 82$ ). All ensembles used in this table have 250 snapshots. See **Methods** for a definition of the evaluation scores.
<sup>a</sup>Default aSAMc model, with 100 reverse diffusion steps.
<sup>b</sup>aSAMc model with no energy minimization.
<sup>c</sup>aSAMc model running with 50 reverse diffusion steps.
<sup>d</sup>Restrained coarse-grained COCOMO simulations followed by all-atom reconstruction via the cg2all neural network. Represents a baseline method.
<sup>e</sup>Ensemble with additional snapshots from the same MD data of the test set ensembles; represents perfect predictions, in the limit of finite-size ensembles.
<sup>f</sup>mdCATH-based aSAMt version. Ensembles were generated at 300 K.
<sup>‡</sup>The scores of these methods were compared with corresponding aSAMc scores (first row) via a two-sided Wilcoxon signed-rank test, which evaluates whether the distribution of differences in paired scores is symmetric around zero.
\*p-value $< 0.05$ in a Wilcoxon test with aSAMc.

We evaluated distributions of Cβ positions using the WASCO-global score^34^, a method to compare protein structural ensembles. Again, AlphaFlow demonstrates a small but statistically significant advantage over aSAMc (**Table 1**). In terms of Cα RMSF correlations and WASCO-global, we note that aSAMc is on the level of ESMFlow, a model trained similarly to AlphaFlow but utilizing the less powerful sequence-based ESMFold network^35^. In any case, all ML methods are largely better than a baseline ensemble generation strategy consisting of coarse-grained simulations^36,37^ with simple elastic network restraints (COCOMO in **Table 1**).

Even though aSAMc and AlphaFlow outperform a simple control, they both struggle at sampling conformations far from the initial MD structure. This is reflected in the mean initRMSD values from ML, which are often lower than the MD ones (**Fig. 2g**). Additionally, WASCO-global scores of the ML methods are largely worse than those of ensembles containing additional snapshots from the MD data of the test systems, which represent optimal ensemble predictions (labeled ‘MD’ in **Table 1**).

We then proceeded to assess the capability to capture backbone torsion angles using WASCO-local scores, that evaluate joint *n* distributions. In this case, aSAMc provides better approximations than AlphaFlow (**Fig. 2c** and **Supplementary Fig. 3a**). This is because, AlphaFlow cannot effectively learn *n* distributions, since its generative modeling training is based only on Cβ positions^18^. Via its latent diffusion strategy, aSAMc can instead learn such distributions. This applies also to side chain torsion angles, where aSAMc approximates much more closely the *χ* distributions from MD (**Fig. 2d**). For all amino acids, *χ* angle pairs are better captured by aSAMc, as exemplified for the 7p41_D test protein (**Supplementary Fig. 3b** and 4). According to our chiJSD score (**Methods**), aSAMc quantitatively outperforms AlphaFlow in modeling such features.

The physical integrity of aSAMc conformations is overall comparable to AlphaFlow, with few heavy clashes and peptide bond length violations (**Table 1**). However, in terms of MolProbity scores^38^, a popular stereochemical assessment method, AlphaFlow outperforms aSAMc by an average of 0.63 units (**Supplementary Fig. 5**). This difference arises because the heuristic energy minimization of aSAMc resolves heavy clashes (severe atomic overlaps), but not soft clashes (minor overlaps) which are also contributing to MolProbity scores. Moreover, energy minimization is necessary for aSAMc to obtain acceptable numbers of heavy clashes (see “SAM no-min” row in the table), while AlphaFlow is minimization-free. This indicates room for improvement in aSAMc.

Finally, in terms of computationally efficiency, aSAMc is faster than AlphaFlow when running both on a GPU (**Supplementary Fig. 6**). Thanks to the lighter networks of aSAMc and its fast minimization protocol (**Methods**), speedups range from 17.4× to 28.0× for proteins with 51 and 554 residues, respectively. The sampling efficiency of AlphaFlow relative to MD simulations from ATLAS has been reported to be at least 10 times higher, we refer readers to the original publication for details^18^. aSAMc speed can be accelerated by decreasing the number of diffusion steps at inference time, the default is 100. When using 50 steps, we measured speedups from 29.0× to 54.4× over AlphaFlow (**Supplementary Fig. 6**), with a negligible decrease in modeling performance (**Table 1**). There is a large deterioration only below 25 steps (**Supplementary Table 2**). Overall, we note that the efficiency of aSAMc is similar to AlphaFlow-Lit, a faster AlphaFlow version with slightly reduced performance^23^.

Taken together, this data shows that aSAMc is comparable to other openly available state-of-the-art models for MD ensemble generation.

### Using aSAMt to learn temperature-dependent behavior from mdCATH

After validating the aSAM architecture against AlphaFlow, we sought to challenge it with a novel type of generation task. Thanks to the mdCATH dataset^21^, we trained a new version, aSAMt, as a temperature-dependent ensemble generator providing temperature as input to its diffusion model. mdCATH has also more sampling with respect to ATLAS. The cumulative simulation time of the training data we used is approximately 55.0 and 0.35 ms for the two datasets, respectively.

We evaluated aSAMt on 90 test set domains from mdCATH which we excluded from training and having at most 20% sequence identity with any training sequence. For such test domains, aSAMt often replicates with low errors several MD average properties and their temperature-dependent behavior. These include initRMSD, radius-of-gyration (*R*_g_), and secondary structure element preservation (SSEP). The first two of these features increase with temperature, while the third decreases with temperature. aSAMt captures such trends (**Fig. 3a** to **c**).

**Fig. 3.**
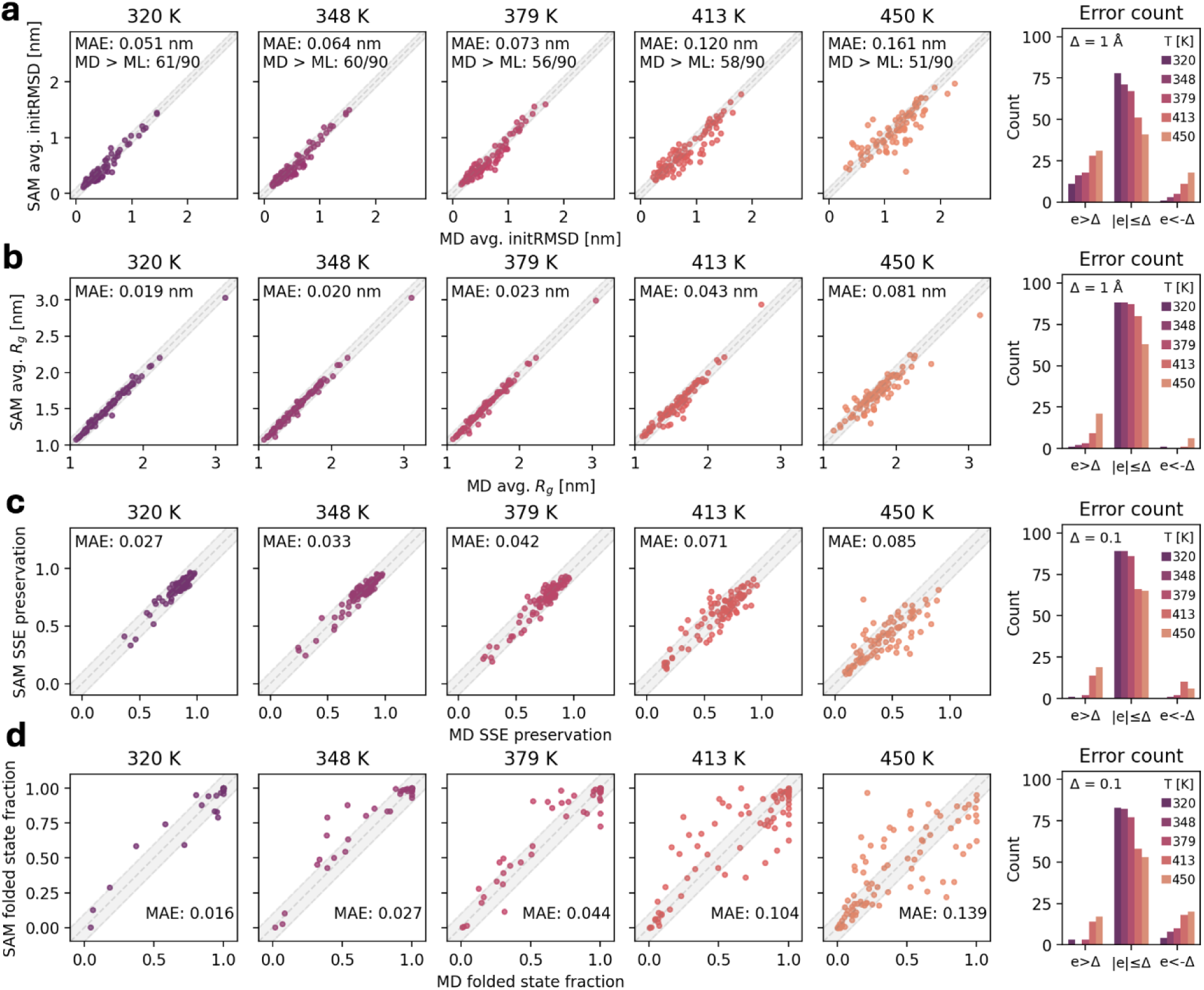
aSAMt approximations of ensemble average properties of mdCATH test set domains. **a** Average initRMSD of MD and aSAMt ensembles for all test set domains (*n* = 90) across five different temperatures (from 320 to 450 K). **b** Average *R*_g_ of the ensembles. **c** Secondary structure element preservation in the ensembles. **d** Fraction of snapshots in the folded state. In each scatterplot (**a** to **d**), we report the mean absolute error (MAE) of aSAMt values relative to the MD ones. For initRMSD value, we also report the number of values underestimated by aSAMt. In the last column, we show the distribution of aSAMt errors. An error *e* is defined as the difference between an MD and an aSAMt value. For each property, we define a threshold Δ (1 Å in **a** and **b**, 0.1 in **c** and **d**) and count errors within or exceeding that threshold. In the scatterplots, aSAMt predictions within Δ correspond to points in the gray-shaded areas. All data are from ensembles of 1250 snapshots.

Interestingly, the capability to replicate average initRMSD values on mdCATH ensembles at 320 K appears better than what the ATLAS-based aSAMc achieves for its test set (**Fig. 2g**). The two models obtain a mean absolute error (MAE) of 0.051 (aSAMt) and 0.108 nm (aSAMc) on their test sets. Additionally, they underestimate average initRSMD in 67.8% (aSAMt) vs. 91.4% (aSAMc) of the cases, showing that aSAMt is more independent from the input structure and achieves broader sampling. Consistent with broader sampling are larger deviations from the ATLAS ensembles for the aSAMc test set when generating such ensembles using aSAMt (**Table 1**). This is reflected in lower RMSF correlations and higher WASCO-global scores for the aSAMt ensembles. We believe that the broader sampling likely results from different sizes of the training sets of otherwise essentially identical models. However, local structural differences in backbone and side chain torsions between the aSAMc and aSAMt generated ensembles (WASCO-local and chiJSD scores in Table 1) likely reflect differences in the force fields used for generating the training data for ATLAS (CHARMM36m) and mdCATH (CHARMM22*).

Protein unfolding is a biophysically-relevant process that can be studied via MD simulations at high temperatures^27,39^. In mdCATH, several domains undergo unfolding. To evaluate the ability of aSAMt to capture such behavior, we used a definition of folded states based on native contacts^40^. We define the folded state fraction (FSF) as the fraction of snapshots in an ensemble that are in the folded state. At 320 K, aSAMt accurately estimates FSFs with a MAE of 0.016 with MD (**Fig. 3d**). However, its performance degrades at higher temperatures and at 450 K it reaches a maximum MAE of 0.141. Despite this trend, aSAMt still qualitatively captures temperature-dependent unfolding for most test domains. For 53 of 90 domains, FSFs from aSAMt at 450 K approximate the MD ones with an absolute error within 0.1. Additionally, aSAMt predicts with an absolute error within 0.1 the FSFs of 23 domains across all five temperatures (**Supplementary Fig. 7**).

To further highlight the ability of aSAMt to capture temperature-dependent conformational sampling, we provide as an example the 4qbuA03 test domain, for which the model recapitulates several MD temperature-dependent properties. This domain is stable at lower temperatures, but at 450 K it starts to unfold. aSAMt replicates the corresponding change in conformational sampling. This can be seen by RMSF values that increment with temperature (**Fig. 4a**) and the unfolding of the two α helices (**Fig. 4b**). In the MD ensemble at 450 K, the domain samples both an unfolded and folded state, as shown by the two-peaked histogram of fraction of native contacts *Q* (**Methods**). aSAMt captures this behavior, reproducing both peaks of *Q* at 450 K (**Fig. 4c**) and FSFs (based on the distribution of *Q*) are closely approximated at all temperatures (**Fig. 4f**). The extent of conformational sampling with aSAMt is also similar to MD (**Fig. 4g**), as can be seen by the temperature-dependent contact maps (**Fig. 4d**) and from the PCA projections of the 3D structures (**Fig. 4e**).

**Fig. 4.**
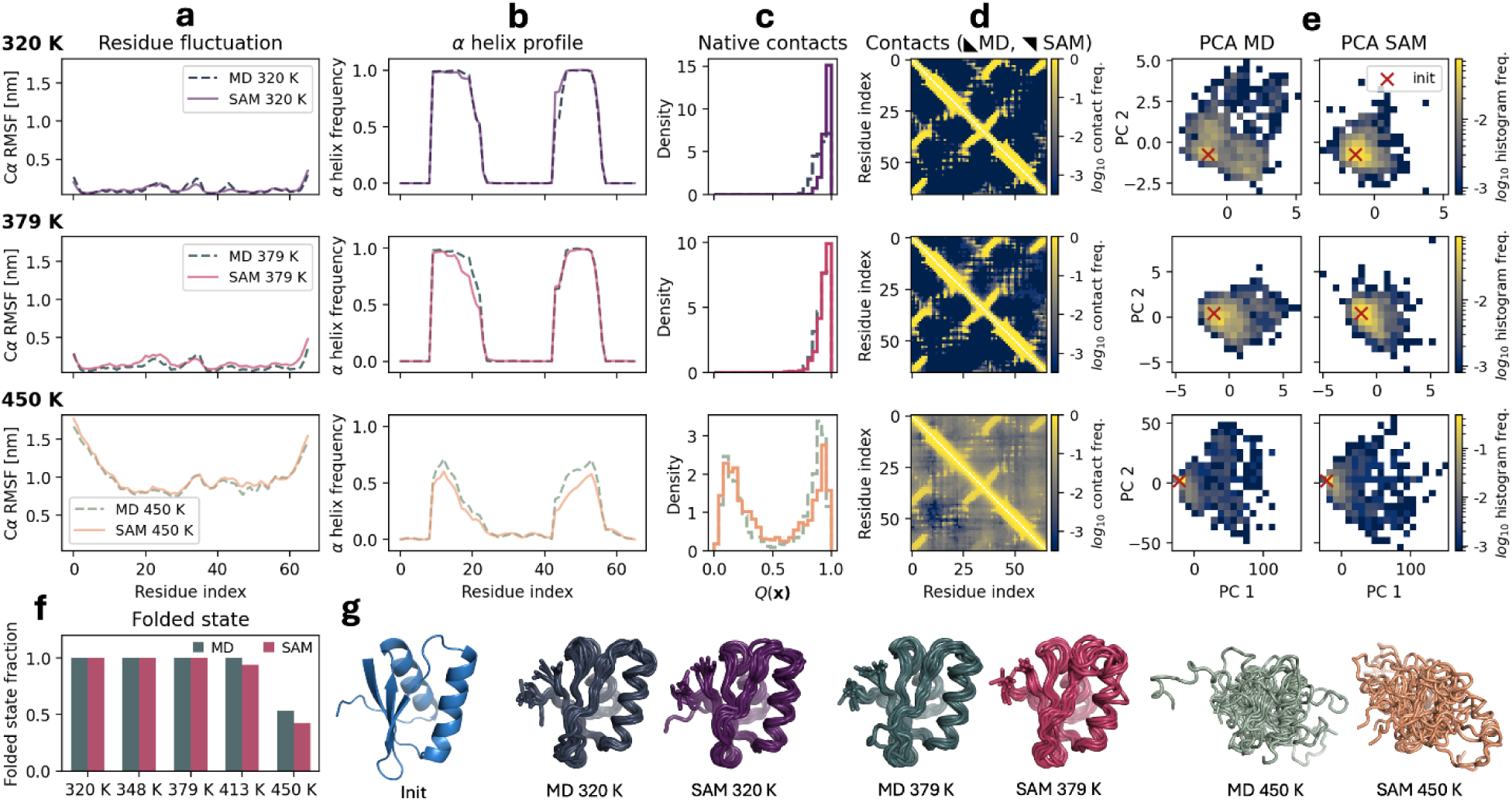
Properties of the ensembles of the 4qbuA03 test set domain from mdCATH. **a** Cα RMSF values of MD (dashed lines) and aSAMt (continuous lines) ensembles of are shown across three increasing temperatures (320, 379 and 450 K). **b** Helicity profiles of the ensembles. **c** Fraction of native contacts *Q* in the snapshots of an ensemble measuring their structural divergence with respect to the initial folded structure of the MD simulation. **d** Cα-Cα contact maps (with a threshold of 0.9 nm) of MD (lower triangles) and aSAMt (upper triangles) ensembles. The color bars report the log_10_ of the contact frequency between residue*i* and *j* in an ensemble. **e** PCA of the ensembles. The red marker represents the location of the initial MD structure. The color bars report the log_10_ of raw histogram frequencies. **f** FSFs of MD and aSAMt across all 5 mdCATH temperatures. **g** Structures of the initial MD conformation (blue) and MD (pale blue shade) and aSAMt (red shade) ensembles. **a** to **g**: All data are from ensembles of 1250 snapshots, the ensembles in **g** are subsamples of 10 structures.

Other similar examples of unfolding induced by temperature are the test domains 2vy2A00 and 3nb2A04 (**Supplementary Fig. 8**). In the MD ensemble at 450 K, these two globular helical proteins are almost completely unfolded and aSAMt can correctly capture this behavior, as reflected in different ensemble properties.

The test set also contains domains that remain substantially folded in MD even at 450 K, like 1qwjB00 and 4a57A02 (**Supplementary Fig. 9**). These large domains of 229 and 366 residues have high *Q* and FSF values at 450 K. 1qwjB00 has a long and flexible loop and the model captures most of its dynamics. For these two domains, even though the conformational space projected via PCA is not perfectly covered by aSAMt, the model can precisely estimate their stability across temperatures, as well as their residue fluctuations and changes in secondary structure content.

Other test set domains already lose stability at 320 K in MD (perhaps because these CATH domains were cropped from larger structures). Interestingly, aSAMt also models this behavior as illustrated by the example 3nb0A03 (**Supplementary Fig. 10**). aSAMt approximates the low folded fractions and secondary structure content at 320 K. At higher temperatures, the MD ensembles become increasingly unstructured and aSAMt again replicates the behavior.

We also provide examples where the ensembles generated by aSAMt deviate from MD. This mostly happens at high temperatures, especially at 450 K. Across the whole test set, aSAMt overestimates (error > 0.1) FSFs for 17 domains and underestimates it (error < -0.1) for 20 domains (**Fig. 3d**). Examples of these two types are shown for 3u28C00 and 3zrhA01 (**Supplementary Fig. 11**). Errors at 450 K are evident from RMSF and secondary structure profiles, where the shapes of the aSAMt profiles generally match the MD ones, but they are shifted to higher or lower dynamics and structure preservation. The errors are also clear from histograms of fraction of native contacts, where the peaks of MD and aSAMt distributions at 450 K are widely separated.

Finally, we evaluated the computational efficiency of aSAMt relative to MD. Using a protocol comparable to the mdCATH one, we simulated via OpenMM^41^ the shortest (4qbuA03) and longest (4a57A02) domains described above. We then compared the GPU times to achieve roughly similar amounts of sampling with MD and ML: 2,500 ns for MD (the mdCATH per-system maximum) and generating with aSAMt an ensemble with 500 snapshots (the maximum snapshot number in mdCATH trajectories, which we deemed sufficient to represent an mdCATH ensemble). aSAMt was 2,317× and 1,000× faster for 4qbuA03 and 4a57A02 (**Supplementary Table 3**), confirming the potential of ML for accelerating MD.

### Interpolation and extrapolation across temperature

Since temperature is provided as a continuous value to aSAMt, the model can sample at any temperature, including those not observed in training. To test aSAMt at a wider range of temperatures, we generated ensembles from 250 to 670 K for the 8 test systems discussed above. We then analyzed their FSFs and SSEPs (**Fig. 5**) to determine the level of folded structures in the ensembles. Both metrics were evaluated to check modeling consistency. For example, the model might erroneously lose native contacts in an ensemble but retain its secondary structure, or vice versa.

**Fig. 5.**
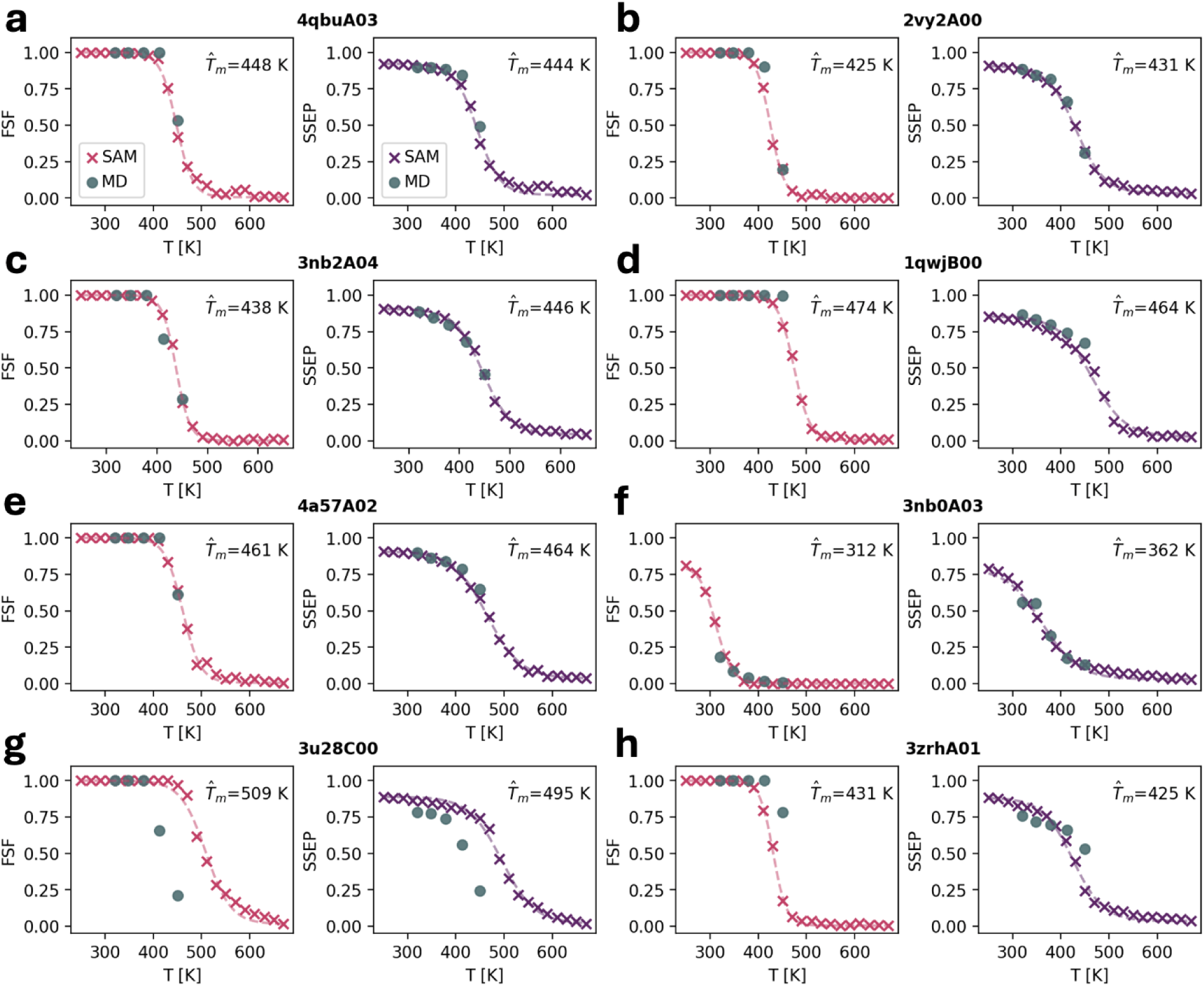
Scanning across temperatures with aSAMt. For 8 selected mdCATH test domains, we generated aSAMt ensembles at temperatures from 250 to 670 K in intervals of 20 K. The folded state fraction (FSF, left subpanels) and secondary structure element preservation (SSEP, right subpanels) are plotted as a function of temperature (red and purple ×). For comparison, we also plot the properties of the MD ensembles of the domains at the five temperatures (320, 348, 379, 413 and 450 K) of mdCATH (gray circles). The dashed curves are sigmoids fitted on aSAMt data. The apparent melting temperatures *T̂*_m_ obtained from such fits are shown for both FSF and SSEP. Ensembles from MD and aSAMt have 1250 snapshots and 200, respectively. **a**-**c** 4qbuA03, 2vy2A00 and 3nb2A04 (partial unfolding in MD), **d**-**e** 1qwjB00 and 4a57A02 (more stable in MD), **f** 3nb0A03 (unstructured in MD), **g**-**h** 3u28C00 and 3zrhA01 (modeling errors from aSAMt).

We found that aSAMt appears to model ensembles at temperatures outside the training data in a plausible way for both FSF and SSEPs. The predicted values interpolate smoothly within the training range and lie close to MD values at temperatures for which mdCATH simulations are available. This interpolation shows that the model has not simply memorized MD behavior at the mdCATH temperatures, but it can approximate temperature in a continuous way.

Notably, values outside the training range also follow physically plausible trends. For most systems, when temperature becomes lower than 320 K, the level of folded conformations remains close to 1 (e.g.: **Fig. 5b**). For the domain already unfolded at 320 K in MD (**Fig. 5e**), aSAMt increases the level of folded conformations when temperature drops, as expected in experiment and from an MD simulation. The model also extrapolates meaningfully at temperatures above 450 K. With increasing temperatures, the level of folded conformations decreases and remains close to 0 for most systems, generating sigmoid-like profiles reminiscent of melting curves observed in experiments^42^ and simulations^43^ of proteins. Even for a system that aSAMt predicted as overly folded at 450K (**Fig. 5g**), unfolded conformations are produced at higher temperatures.

The data from aSAMt allows us to fit sigmoids (dashed curves in **Fig. 5**) and obtain apparent melting temperatures *T̂*_m_. The *T̂*_m_ values obtained with FSF or SSEP are typically similar, confirming the consistency of the model.

We further illustrate ensemble properties across the whole range of temperatures for 1qwjB00 (**Supplementary Fig. 12**). As temperature increases, secondary structure elements unfold and RMSF amplifies, with a more pronounced change near the *T̂*_m_. aSAMt ensembles retain residual secondary structure at high temperatures, consistent with experimental observations of residual helicity in denatured proteins under various conditions^44^. However, since the model was trained on high-temperature trajectories started from folded structures, some residual structure may also be expected simply based on the training data.

To further check the performance of aSAMt with out-of-training temperatures, we monitored additional properties (**Supplementary Fig. 13**). The average initRMSD of the domains behaves in a consistent way, with lower fluctuations below 320 K and increasing divergence at higher temperatures. The average *R*_g_ remains constant up to the *T̂*_m_ of a system and then begins to increase as the domain unfolds, but stays below the *R*_g_ of a non-interacting ideal polymer^45^, consistent with a moderate level of compaction expected in unfolded ensembles of proteins^46^. The number of heavy clashes tends to increase with temperature despite energy minimization, but it stays lower than 1 per molecule for most systems at 670 K. Since we developed aSAMt for lower temperatures, we leave a solution to this problem for future studies.

### Energy landscapes of fast-folding proteins

An ideal target of ML ensemble generators is to explore protein energy landscapes, discover metastable states, and help characterize folding principles, because these tasks are computationally prohibitive for MD. We asked to what extent the mdCATH-based aSAMt could meet this challenge, at least for small proteins. Specifically, we focused on the question whether aSAMt can explore the full landscape of such proteins at higher temperatures. To answer this question, we compared aSAMt ensembles with long unbiased MD simulations of 12 fast folding proteins performed on the ANTON special-purpose hardware at D.E. Shaw Research^12^ (**Supplementary Table 4**). In this dataset, proteins undergo multiple folding and unfolding events and exhaustively explore their landscapes. For each protein, we generated aSAMt ensembles at the same temperature of the MD simulations (in the 290-370 K range) and at 450 K, each with 500,000 conformations. The input of aSAMt is the reference PDB structure of a protein. We directly used the mdCATH-based aSAMt, with no re-training, since the long MD simulations used for comparison were performed with the same CHARMM22* force field^12^ used for the mdCATH training data.

Ensembles generated by aSAMt were projected onto coordinates from time-lagged independent component analysis^11^ (TICA) based on the MD energy landscapes (**Fig. 6**). In all aSAMt ensembles at the given MD temperatures the free energy minima corresponded to the native state as in the MD (black markers). However, for most systems aSAMt also extensively sampled non-native conformations, including non-native basins, similar to the MD simulations. Despite good qualitative agreement, free energy values from aSAMt were not in accurate quantitative agreement with the long MD results indicating that aSAMt learned the extent of conformational sampling covering a broad range of conformations but struggles in correctly reproducing relative probabilities between different states . This result may be expected, since the training data from mdCATH were simulations at most 500 ns long with few transitions between alternate states at a given temperature from which relative probabilities could have been learned. Interestingly, for several systems, the exploration of non-native unfolded states is augmented when the temperature is increased to 450 K, as seen for the WW-domain, NTL9, BBL and the homeodomain. In aSAMt ensembles at 450 K, the free energy of such states is typically lower than for the native state, reflecting the expected effect of thermal unfolding. However, for most of the proteins the MD ensembles include additional sampling of conformations that are separated by a large kinetic barrier from other states, which are not reached by aSAMt even at 450 K. Examples include the WW-domain, NTL9, BBL, protein B, alpha3D and λ-repressor.

**Fig. 6.**
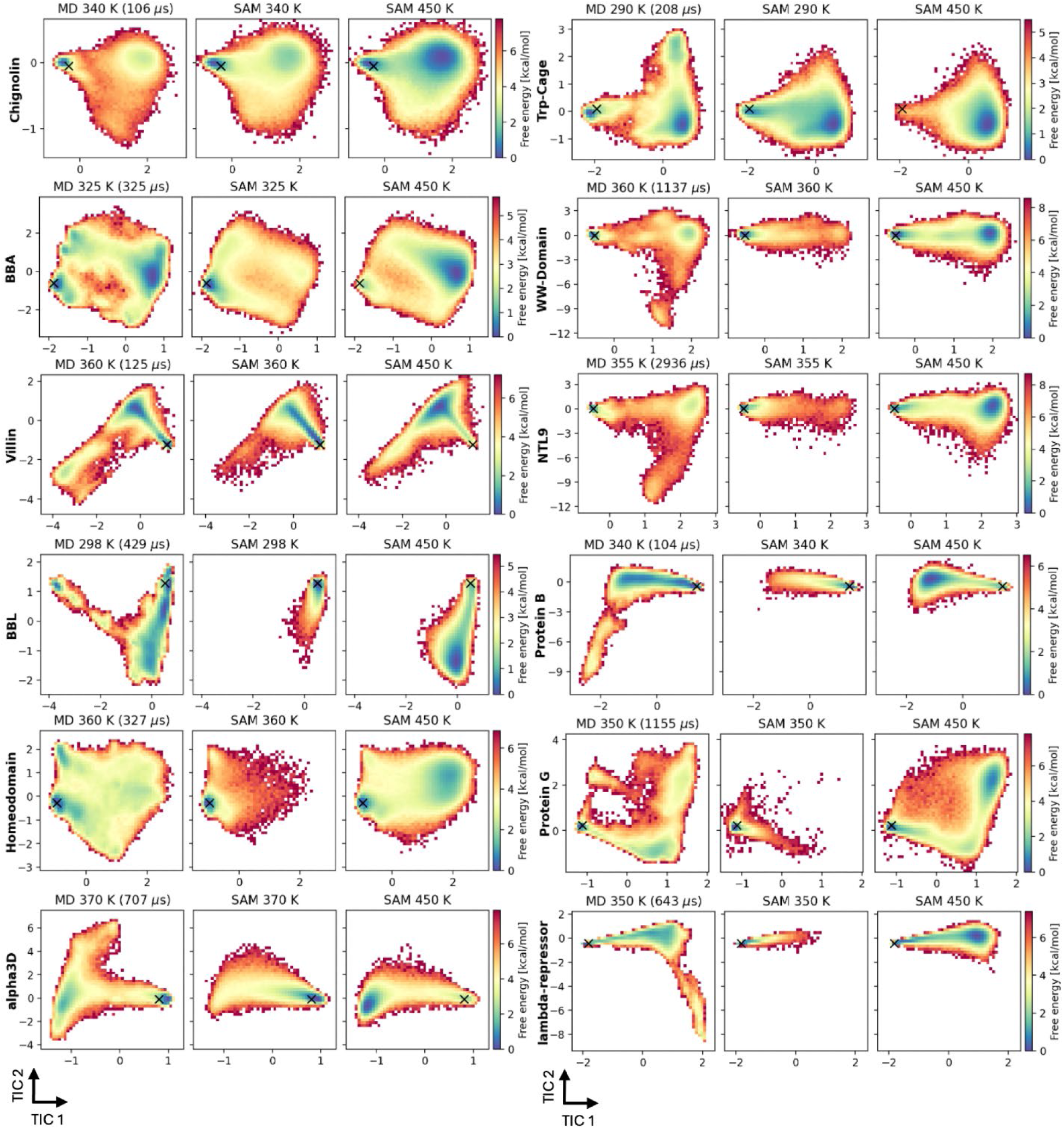
Energy landscapes from long MD simulations and aSAMt ensembles. MD simulations of 12 fast folding proteins performed at D.E. Shaw Research were used to build free energy landscapes via TICA dimensionality reduction. The mdCATH-based aSAMt ensembles for the same proteins were projected on such landscapes. For each protein, we show the MD data (left panels) the aSAMt ensemble generated at the MD temperature (center) and a aSAMt ensemble at 450 K (right). We used 500k snapshots randomly extracted from MD data or generated by aSAMt for each panel. The simulation time of the MD trajectories is shown in parenthesis. A black × represents the location of the input structure of aSAMt. In all subplots, the horizontal and vertical axes represent the first two time-independent components (TIC). Color bars report free energies.

To check whether the aSAMt ensembles for the fast-folding proteins may have been examples of training set memorization, we searched for structural similarity of each protein in the mdCATH training set based on the highest TM-score^47,48^. For six proteins we found a homologous mdCATH domain (**Supplementary Fig. 14**), with NTL9 and lambda-repressor having matches with a 100% and 96% identity over aligned regions. Interestingly, when we projected training snapshots onto long MD landscapes, we observed that aSAMt generates conformations in regions absent in mdCATH (**Supplementary Fig. 15**). For example, for protein G at 450 K, the model explores non-native basins barely sampled in mdCATH. In general, aSAMt ensembles at the temperature of the long MD simulation cover more regions than the training data at the closest temperature. Therefore, aSAMt does not merely memorize training snapshots, rather it interpolates, and moderately extrapolates the in generated conformational space. However, it is clear that significant extrapolation to non-native states that are kinetically far from the native state remains a key challenge, as for other transferable ML models^26^ and even large scale MD efforts^49^.

### Estimation of thermostability and comparison with experiment

Given the ability of aSAMt to explore conformational spaces across temperatures, we next asked to what extent the model could approximate experimentally observed thermostability of proteins. The mdCATH data consists of MD simulations with a force field not explicitly parametrized to reproduce the effect of temperature on protein stability^50^. Consequently, quantitative agreement with experiment cannot be expected from a model trained on such data. However, we thought that even qualitative agreement from a fast ML ensemble generator trained on simulations might constitute a step towards future models incorporating experimental data in training^26,51^.

We first investigated whether aSAMt could capture the trends in experimentally measured *T*_m_values. We utilized a dataset from Pucci et al.^52^, containing a list of PDB entries of proteins with an experimental *T*_m_. From this dataset, we selected 62 monomeric proteins with no cofactors or disulfide bridges to stabilize the structures (**Methods**). This was necessary because aSAMt was not trained to consider such features, which play a crucial role in protein stability, are absent in mdCATH simulations. We then generated aSAMt melting curves (**Supplementary Table 5**) based on folded state fractions (**Fig. 4a**) and obtained *T̂*_m_ values to compare with the experimental values (**Fig. 7a**).

**Fig. 7.**
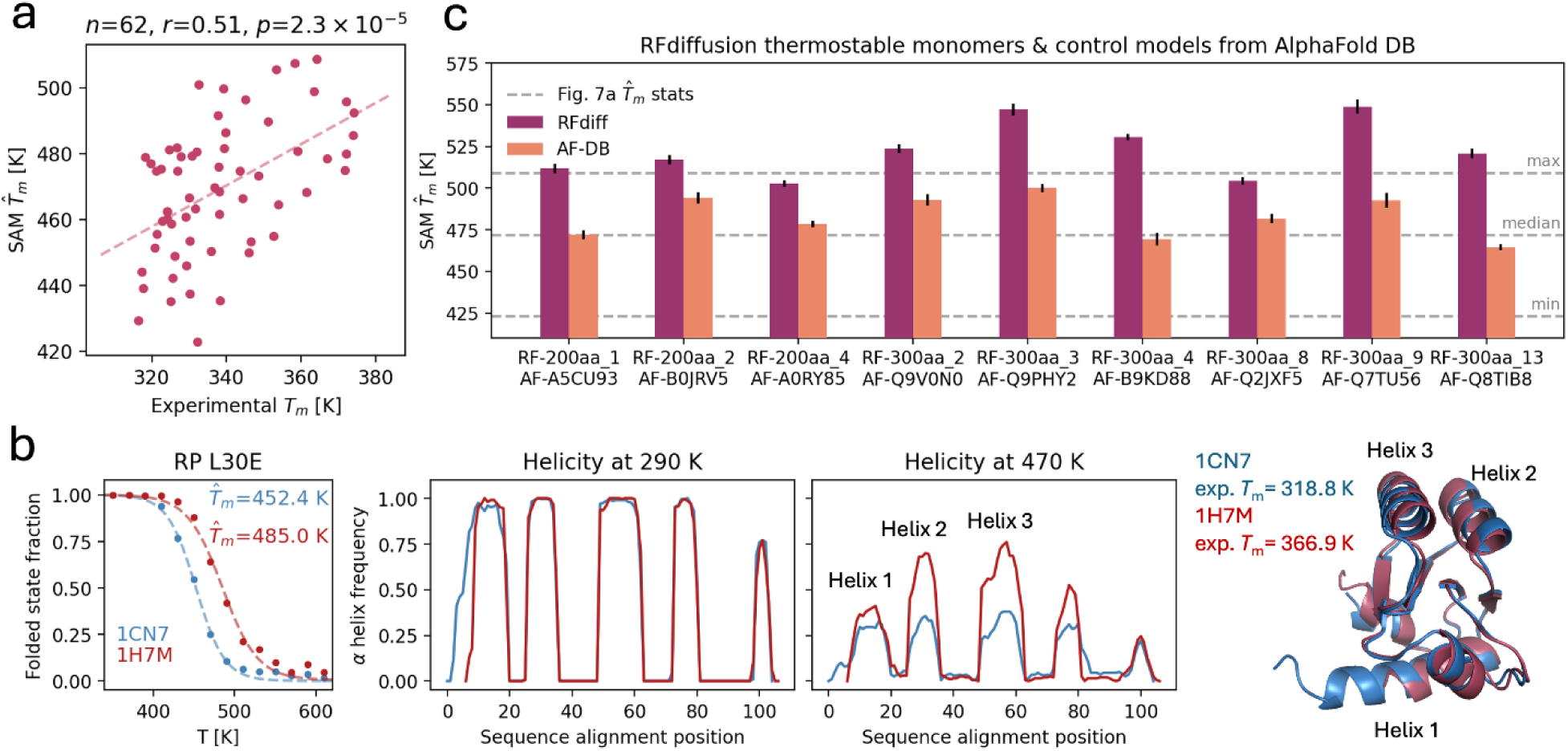
Modeling proteins with experimentally observed thermal behavior. **a** Experimental *T*_m_ values compared with *T̂*_m_ values from aSAMt for a set of monomeric proteins (*n* = 62). The figure shows a regression line (dashed), the Pearson correlation coefficient *r* between experimental and aSAMt values and its associated two-sided *p*-value, calculated via the exact density function for the sample correlation coefficients of a normal bivariate. **b** In the right subpanel, aSAMt melting curves are shown for a pair of ribosomal L30E proteins. The *T̂*_m_ values for the mesophile protein (PDB 1CN7, blue) and the hyperthermophile (PDB 1H7M, red) are also reported. In the central subpanels, the α helix profiles in aSAMt ensembles at 290 and 470 K are shown. Peaks in the profiles correspond to α helices in the native 3D structures (shown on the right). Experimental *T*_m_ values are shown beside. **c *T̂***_m_ values for 9 monomeric proteins designed by RFdiffusion (red) are compared to *T̂*_m_ values of AF2 structures with matching structural properties (orange). The horizontal gray lines show the minimum, median and maximum *T̂*_m_ values for the proteins in panel **a**. The bars report the mean and standard deviation for 5 *T̂*_m_ values obtained from replicas of 50 snapshots. **a**-**c**: details on the number of snapshots and temperatures ranges used of aSAMt are reported in **Supplementary Table 5**.

The values from aSAMt are systematically 130 K higher than experiments. We believe that this might be caused by force-field limitations as well as limited-length training simulations that start from a folded structure and therefore overestimate foldedness, and aSAMt errors at replicating MD behavior at higher temperatures (**Fig. 3d**). Despite this, *T̂*_m_ values estimated from aSAMt ensembles are correlated in a significant way (*r*=0.510, *p*=2.3×10^-5^) with experimental data. We note that the model was not explicitly trained as a *T*_m_ predictor and to obtain estimates of *T̂*_m_ it was often necessary to extrapolate significantly beyond training temperatures (**Fig. 5**). The fact that it generated zero-shot *T̂*_m_ values roughly correlated with experiments confirms that it learned, to at least some extent, a meaningful approximation of temperature effects on the conformational ensembles of proteins.

Next, we analyzed homologous proteins with different experimental *T*_m_ to explore whether a generative model can probe sequence properties influencing thermal stability^53^. We examined five protein pairs^54^ with characteristics similar to those indicated above (**Supplementary Table 6**). Here, we highlight two ribosomal L30E proteins from a mesophile and a hyperthermophile^55^ with an experimental Δ*T*_m_ of 48.1 K. aSAMt qualitatively captures this difference producing a Δ*T̂*_m_ of 32.6 K. The model suggests a hypothesis for such behavior, as in its ensembles two of the five α helices in the hyperthermophile protein exhibit increased stability at high temperatures (**Fig. 7c**).

For other homologous pairs, aSAMt replicates the sign of experimental Δ*T*_m_ (**Supplementary Fig. 16**). However, for the two pairs with sequence identity above 50%, it produces curves with relatively small Δ*T̂*_m_. This limitation may stem from training data, which includes proteins with at most 20% identity, perhaps hindering the ability to model effects of more subtle sequence variations.

Finally, we explored whether aSAMt could also model *in silico* designed proteins^56^. Such proteins are relevant in biomedical research and are good transferability tests for ML models. We analyzed 9 proteins generated by the RFdiffusion generative model which, when expressed, were characterized as extremely thermostable^57^. Since their experimental 3D structures are not available, we used AF2 models instead as input to aSAMt. The *T̂*_m_ values for 7 of these proteins (**Fig 7c**) were higher than any *T̂*_m_ of the natural proteins (**Fig 7a**). To check if the cause was the use of AF2 input, we performed a control utilizing 9 models of natural proteins from AlphaFold Database^58^, selected to have the same length and the highest possible structural compatibility to the designed proteins (**Methods**). *T̂*_m_ values for such controls were always lower than the RFdiffusion counterparts (**Fig 7c** and **Supplementary Fig. 17**), showing that aSAMt can specifically capture thermostability in proteins designed by deep learning.

## DISCUSSION

Early deep generative models for protein ensembles were system-specific models with limited applicability beyond their training systems^59^. Lately, more transferable models have emerged, holding promise for broader biomolecular dynamics studies, with idpGAN^60^, idpSAM^32^, AlphaFlow^18^ and, most recently, BioEmu^26^ as pioneering examples. Here, we propose a novel MD- based ensemble generator that further advances the field by achieving higher accuracy in modeling side chain and backbone torsion angles and, for the first time, allowing to incorporate temperature effects in the generative process.

Our model aSAM is an SE(3)-invariant latent diffusion model, derived from idpSAM^32^. Latent generative models of protein structures were recently explored in other studies, such as Structure Language Modeling^61^ and LATENTDOCK^62^. As many ensemble generators operate on Cartesian coordinates and frames^3,18,25,26^, it is useful to develop and compare different solutions. For example, aSAM can efficiently model side chain distributions, contrary to most other models. Since every atom is encoded in the latent space, a latent model can naturally learn their joint distributions. A robust decoder, like the AF2 Structure Module, can then map latent representations to 3D structures respecting the learned distributions. While aSAMt generates only heavy atoms, it could be easily extended to hydrogen atoms^37^, to enable applications that require hydrogens such as the integration of NMR restraints^51^. Intriguingly, while we trained two AEs for ATLAS and mdCATH, we observed good cross-dataset performance (**Supplementary Table 1**). This hints that a unified AE model of protein structures could serve as a foundation for future latent generators, requiring only a generative model component to be trained when using new datasets (**Fig. 1b**). AEs of this kind have already been explored^63,64^.

In terms of modeling performance on the ATLAS dataset, the ATLAS-based aSAMc is comparable with open models like AlphaFlow. Our model captures backbone and side chain torsion angle distributions with considerably higher accuracy. However, it has slightly inferior performance on positional distributions and residue fluctuations (**Table 1**). We attribute this to two factors. First, AlphaFlow is fine-tuned from AF2, which was pre-trained on the PDB, and probably has a richer prior understanding of protein structures and their variability. aSAMc was instead directly trained only on part of the ATLAS training set. Secondly, AlphaFlow uses AF2-derived attention mechanisms likely possessing stronger inductive biases compared to the regular self-attention adopted in aSAMc, possibly improving its data efficiency. These factors could lead to a statistically significant better generalization on test data.

Because of the limited size ATLAS, both the ATLAS-based aSAMc and AlphaFlow struggle to sample states distant from initial MD structures. In contrast, we found improved performance for the mdCATH-based aSAMt compared to aSAMc in capturing MD variability. We attribute this result to the larger size of the mdCATH dataset. The recently introduced BioEmu appears to also overcome this challenge and captures multiple states in large proteins. Given that BioEmu has a similar generative model and neural network architectures with respect to aSAM and AlphaFlow, we believe that BioEmu behavior likely stems from its different training strategy and a larger training set. We remark that our study was made possible by large, open MD datasets like ATLAS and mdCATH. As current generative models are essentially data-driven statistical machines, development of open ML ensemble generators is dependent on the availability of such datasets.

aSAMt is one of the first examples^65^ that show how transferable generators trained on limited sampling data can explore, albeit with incorrect probabilities, large portions of the full energy landscapes of small proteins, such as the 12 fast folding proteins analyzed here^12^. The combination of numerous training systems and high-temperature simulations, like in mdCATH, is likely at the core of this capability. Interestingly, a BioEmu version has also been trained in a one hold out fashion on the same D.E. Shaw simulations of fast folding proteins that we considered (**Fig. 6**). BioEmu ensembles are not currently available, but from the published results, it appears that the ensembles generated by aSAMt are similar to those from BioEmu^26^, even though aSAMt was not retrained on the fast folding proteins long MD simulations in a one-hold out fashion like BioEmu. Notably, both aSAMt and BioEmu appear to miss the same kinetically distant states (for WW-domain, NTL9, BBL, Protein B, α3D and λ-repressor).

Such results confirm that MD-based ensemble generators still face two challenges^17^: learning the correct equilibrium probabilities (i.e.: free energies) from MD and covering all relevant states in a system. A possible strategy to the first problem, is to reweight training snapshots from out-of-equilibrium short MD trajectories with equilibrium weights, obtainable for instance by Markov state modeling^66^ or experimental data^26^. The coverage problem is likely a more fundamental limitation. We note that some of the fast-folding proteins analyzed here were also recently simulated by Majewski et al.^49^. The authors used the same force field from mdCATH and an adaptive sampling strategy with thousands of short unbiased trajectories, accumulating up to ms of sampling per system, which they publicly shared. Yet, when we compared these simulations with D.E. Shaw data, we observed that the states not reached by aSAMt and BioEmu sometimes also remained unexplored in the adaptive MD simulations (**Supplementary Fig. 18**). This may suggest that training with short MD, which may be limited in reaching states that require crossing of multiple, larger energy barriers, may not allow ML generators to learn how to reach such states. An alternative explanation, that is beyond the scope of this work to demonstrate, is that the unbiased simulations from D.E. Shaw may have explored kinetically trapped rare states with overestimated stability due to limited sampling, despite very long simulation times.

A key advance in our work is the incorporation of environmental conditions in ensemble generators. To our knowledge, aSAMt is the first example of a conditional and transferable generative model of this kind. While here we focus on temperature, the approach could be extended to other factors, including other physicochemical variables or experimental restraints on a path towards achieving broader generalization. It is clear, however, that this remains a very ambitious goal to replace physics-based MD techniques with the current generation of deep generative models likely requiring extremely large datasets. However, the newly emerging ML methods, including aSAM, may complement MD as heuristic tools, especially when considering high-throughput studies^67^. Additionally, hybrid strategies that combine the strengths of MD and ML have already been described^68^ and further improvements in ML models will unlock new applications.

A fundamental challenge in ML ensemble generators is bridging the gap between simulations and experiment. This challenge is exacerbated by the fact that sufficiently large training data are only available from simulations, which are themselves limited in reproducing experimental data. Moreover, environmental complexity makes it difficult to accurately reproduce experimental behavior, as seen in our comparison of aSAMt predictions with *T*_m_ values from experimental measurements (**Fig. 7**). While aSAMt qualitatively captures *T*_m_ of several proteins, we excluded proteins with properties absent from training (e.g.: oligomers). We note that many state-of-the-art *T*_m_ predictors are in fact sequence-based^69^, as they implicitly capture biomolecular interactions embedded in sequences. Incorporating experimental data into simulation-based ML models, either to provide a more complete picture of the environment^51^ (e.g.: type of solvent) or to overcome force field and sampling limitations^26^, represents a promising direction to improve experimental agreement and increase the practical relevance of ML ensemble generators.

## METHODS

### ATLAS dataset

To compare aSAMc with AlphaFlow, we used the ATLAS^22^ dataset (https://www.dsimb.inserm.fr/ATLAS/index.html), which consists of explicit solvent simulations of 1,390 proteins chains from the PDB^9^. For each chain, there are three 100 ns simulations. We utilized the same training, validation and test set splits of AlphaFlow^18^. Due to computational constraints, we removed all chains with more than 500 residues in the training and validation sets, resulting in 1,174 and 38 chains, respectively. For the test set, we used all 82 proteins, with lengths from 39 to 724.

### mdCATH dataset

To train a temperature-conditioned aSAMt, we used the mdCATH^21^ dataset (https://github.com/compsciencelab/mdCATH), containing explicit solvent simulations of 5,398 protein domains from CATH^70^. Simulations are available at five temperatures: 320, 348, 379, 413 and 450 K. For each domain and temperature, there are five trajectories of at most 500 ns. We randomly split the domains into 5,268 training, 40 validation and 90 test elements. Since mdCATH domains are already clustered at 20% sequence identity, this split effectively yields partitions that do not share highly similar domains. The maximum chain length of mdCATH is 500. For training the autoencoder and diffusion models, we applied a length cutoff of 320, resulting in 5,056 domains. We did not apply cutoffs for the validation and test sets.

### aSAM model

The aSAM model (referring to both aSAMc and aSAMt) is based on our previously described Cα- based idpSAM^32^ method, with two main differences. First, here we used largely different neural networks in the autoencoder to model all protein heavy atoms. Second, while the diffusion model is similar, its neural network has new modules to embed input conditions and achieve an overall increased capacity.

#### Autoencoder

The first stage of aSAM training consists in learning to encode the positions of protein atoms into SE(3)-invariant representations via an autoencoder (AE). We define an encoder *E*_*φ*_, with learnable parameters *φ*, as:

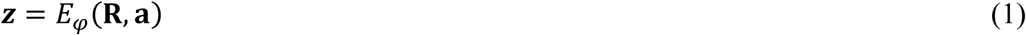

where **R** ∈ ℝ^*Nh*×3^ are the Cartesian coordinates of *N*_*h*_ heavy atoms in a protein with *L* residues, **a** ∈ ℝ^*L*×1^ is its amino acid sequence and **z** ∈ ℝ^*L*×*c*^ is an encoding (a latent representation) for the 3D structure. We aim to learn SE(3)-invariant encodings:

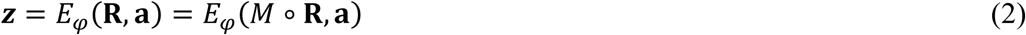

for any transformation *M* with translations or rotations that does not reflect coordinates. The encoder is implemented as a transformer-based network with four blocks. To provide a rich representation of **R**, its inputs include radial basis expansions^71^ of the full distance matrices of Cα- Cα, N-O atoms and SCC-SCC beads (SCC: side chain centroid) and features from *ϕ*, *ψ*, *ω*, *χ*_1_ to *χ*_4_torsion angles and local frame-aligned^1^ Cα and SCC positions. Since these inputs are roto-translationally invariant, encodings inherit the same invariance. Here, we fix the latent dimension to *c* = 32. Each of the *L* tokens in an encoding corresponds to a residue of the original chain, although it likely carries structural information from multiple residues because of the flow of information in the transformer-based encoder.

We next define a decoder *D*_*ξ*_, with parameters *ξ*, as:

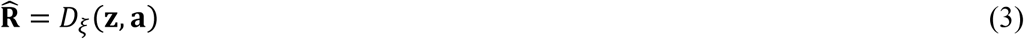

where **R̂** ∈ ℝ^*Nh*×3^ is a reconstructed heavy atom conformation. The decoder is based on the Structure Module from AF2, with five blocks^1,72^. The module is preceded by minimal input layers mapping **z** and **a** to a higher-dimensional space.

The AE objective function is based solely on reconstruction and loss terms capturing stereochemical quality. The loss function for a single conformation is:

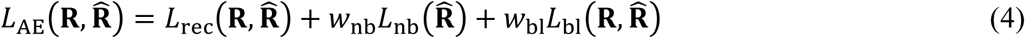

where **R̂** = *D*_*ξ̂*_ *E*_*φ*_(**R**, **a**), **â** is an autoencoded 3D structure, *L*_rec_ is a loss term based on the frame aligned point error (FAPE) and auxiliary losses from AF2, *L*_nb_ is a term penalizing non-bonded Cα atoms from getting too close, and *L*_bl_ enforces the correct distance between adjacent Cα-Cα. *w*_nb_ and *w*_bl_ are weights. Detailed descriptions of the AE neural networks and their training are given in **Supplementary Note 1**, **Supplementary Fig. 19** and **Supplementary Table 7**.

#### Latent diffusion model

Using the encoder of the AE with frozen weights, it is possible to encode a dataset and to apply a generative model to learn the data distribution in the latent space. We choose as a generative model the denoising diffusion probabilistic model (DDPM) formulation of Ho et al.^20^. We used the same DDPM framework as in the idpSAM model^32^.

A DDPM is based on a noise prediction network **ϵ_*θ*_**, with learnable parameters *θ*. Here, the task of **ϵ_*θ*_** is to receive as input a noisy encoding **z**_***t***_ ∈ ℝ^*L*×*c*^, representing a noisy 3D structure of a protein chain, along with the corresponding diffusion time step ***t*** and to output the noise with which **z**_***t***_ was perturbed. During DDPM training, the encoder is set as non-trainable. Our **ϵ_*θ*_** is conditioned also on the amino acid sequence **a** of the chain, the 3D structure used for MD initialization and, for a temperature-dependent model, a scalar value representing temperature. The network has a transformer-like architecture inspired by the Diffusion Transformer^73^, which readily allows to integrate such inputs in the denoising prediction.

We trained the DDPM with the variational lower bound loss *L*_simple_ from the DDPM paper^20^. For the diffusion process, we used a maximum timestep of 1,000 and a sigmoid noise schedule^74^. Details on **ϵ_*θ*_** and its training are in **Supplementary Note 1**, **Supplementary Fig. 20** and **Supplementary Table 8**.

#### Input structures

Depending on the dataset, we use different definitions of initial structures for **ϵ_*θ*_**. We avoided directly using structures from the Protein Data Bank as they may be of low resolution and/or subject to technical issues, such as missing atoms or modified residues.

For the ATLAS-based model, we used the PDB structure that accompanies the MD trajectories of each chain in the database. This is described by the ATLAS authors as the structure used to launch MD simulations after energy minimization and equilibration.

For mdCATH, we use the first MD snapshot from the trajectories at each temperature during training, and at 320K for evaluation on test set domains, irrespective of the temperature at which we sample.

For proteins outside mdCATH, we use initial conformations of varying origins.

#### Sampling

When sampling with the diffusion model, we employ a lower number of reverse diffusion steps with respect to the maximum used in training. Unless otherwise stated, all results here are obtained with 100 steps. For decoding the generated encoded conformations, we directly use the *DD*_*ξξ*_ network as a deterministic decoder^32^.

#### Final minimization

To relax heavy atom clashes, we apply a restrained energy minimization protocol implemented in PyTorch. We include all protein bond length and bond angles terms and selected improper and proper dihedral terms from the Amber ff99SB force field^75^. The restrained properties are backbone *ϕ*, *ψ* torsion angles and side chain *χ*_1_ to *χ*_4_ torsion angles as well as adjacent Cα-Cα distances. We restrain these features to their values in the snapshots generated by aSAM, because this simplified minimization would otherwise deteriorate their distributions. Details of the protocol are given in **Supplementary Note 2**.

### Evaluation of ATLAS ensembles

All evaluations on the ATLAS test set were performed with ensembles of 250 conformations, like in the AlphaFlow publication^18^. In case of reference MD ensembles, snapshots were uniformly sampled from the three available simulations. To establish a positive control for a perfect ensemble prediction, we extracted an additional 250 snapshots from the same MD data and compared it against the first 250 snapshots.

To compare aSAMc with AlphaFlow and ESMFlow, we downloaded the ATLAS test set ensembles with 250 snapshots provided at https://github.com/bjing2016/alphaflow?tab=readme-ov-file#Ensembles. We used the AlphaFlow-MD+Templates and ESMFlow-MD+Templates versions of these ensembles, derived from models that also take as input an initial MD structure.

To compare the ML methods to a simple ensemble generation baseline, we performed restrained coarse-grained MD simulations of the ATLAS test set proteins using the COCOMO2 Cα-based force field^36^ in OpenMM 8.0.0^41^. Each system was minimized and equilibrated for 5,000 steps before running 0.5 µs production simulations, generating 5,000 frames per trajectory. Simulations were performed in the NVT ensemble at 298 K, using Langevin dynamics with a friction coefficient of 0.01 ps⁻¹ and a 10 fs integration time step. Systems were placed in a 30 nm cubic periodic box, with nonbonded interactions modeled using a 10–5 Lennard-Jones potential and a Debye–Hückel potential for electrostatics. Nonbonded interactions were scaled based on solvent exposure to account for solvent effects. To preserve the structure of folded proteins, an elastic network model was applied, with harmonic restraints between Cα atoms within a 0.9 nm cutoff and a force constant of 500 kJ/mol across non-adjacent residue pairs. To obtain the ensembles used in **Table 1**, 250 snapshots were then extracted from the CG trajectories and atomistic details were reconstructed with the cg2all network^37^.

To compare the generated structural ensembles with the reference (MD) ones, we used a series of scores:

#### Root-mean square fluctuations (RMSF)

To evaluate RMSF, we computed the Cα RMSF of the MD and generated ensembles and calculated the Pearson correlation coefficient (PCC) between the two series. RMSF values were computed using MDTraj^76^, with all conformations from a given ensemble superimposed onto the initial structure of the MD simulation.

#### WASCO

To evaluate distributions of 3D positions of Cβ atoms and joint distributions of *n* backbone torsion angles, we utilized the WASCO-global and WASCO-local scores^34^, respectively. These correspond to the “overall global discrepancy” and “overall local discrepancy” values in the WASCO article and are computed as in the *comparison_tool.ipynb* notebook of its package. We used default parameters of the notebook and no ensembles replicas. Both scores are based upon Wasserstein distances between distributions, with values closer to 0 corresponding to better agreement.

#### Side chains

To evaluate distributions of side chain torsion angles *χ*_1_ to *χ*_4_, we developed the chiJSD score:

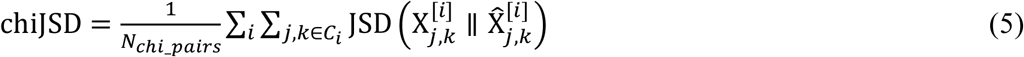

where *i* is the residue index in a chain of length *L* and *C*_*i*_ is the set of existing pairs of *χ* indices of residue *i*. The pairs are (1, 2), (2, 3) and (3, 4), depending on the amino acid type. 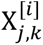 and 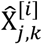 denote the bidimensional distribution of angles *χ*_*j*_ and *χ_k_* for residue*i* in the reference and proposed ensembles, respectively. JSD represents the Jensen-Shannon divergence and *N_chi_pair_* is the number of evaluated distributions. To compute JSDs, we discretized observation into 2D histograms^77^. For both dimensions, we defined the bin range (−120°, 0°, 120) and we wrapped values outside the range onto the same bins around the 2D torus, yielding a total of 9 equally sized bins. We used such bins because they fit the natural distribution of several *χ* angles^78^ (**Supplementary Fig. 3**-4). chiJSD ranges from 0 to ln(2), with lower values corresponding to better agreement between distributions.

#### Physical integrity

To assess the integrity of ML conformations, we evaluated the number of heavy-atom clashes and violations in N-C peptide bond lengths per snapshot. We focused on these properties because aSAMc and AlphaFlow utilize the Structure Module of AF2, which enforces most bond lengths and angles to standard literature values. To define “heavy-atom clashes”, we considered all pairwise distances between non-hydrogen atoms from non-adjacent residues in a conformation. We then counted distances where 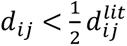, with 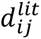 being the sum of van der Waals radii of the atoms. To define “peptide bond length violations”, we counted all peptide bond lengths *b*_*i*_ for which |*b*_*i*_ − *b*_m_|⁄*b*_s_ ≥ 3.0, where *b*_m_ = 1.348 Å and *b*_s_ = 0.029 Å are the mean and standard deviations for N-C lengths in the ATLAS training data. The scores reported in **Table 1** and **Supplementary Fig. 13** are the number of such clashes or violations per snapshot in an ensemble, respectively.

#### Principal component analysis (PCA)

The PCAs in **Fig. 2** and **4** were performed using Cα-Cα distances from all pairs of non-adjacent residues. The analysis was performed on MD data first and the ML ensembles were then projected on the axes derived from the MD sampling. For mdCATH ensembles, we carried out PCA for each temperature.

#### Timing

To measure the speed of aSAMc and Alphaflow, we generated 250 snapshots for 11 ATLAS test proteins with lengths from 51 to 554. For AlphaFlow, we used the *predict.py* script in its package with default settings. We ran both models on NVIDIA V100 32GB GPUs and measured exclusively the time spent for sampling (e.g.: we excluded time for loading weights and constructing MSAs). aSAMc was used to generate samples with a batch size of 1 or 8, and by sampling with 50 or 100 diffusion steps. AlphaFlow uses a batch size of 1, and 10 steps. For SAM, the total time we report is the sum of the time for generating 3D structures and for energy minimization (roughly 5-10% of the process). We used a batch size of 50 for the minimization.

### Evaluation of mdCATH ensembles

All evaluations on mdCATH test set domains were performed with ensembles of 1,250 conformations. For a reference MD ensemble at a given temperature, snapshots were uniformly sampled from the five available simulations. To analyze aSAMt ensembles, we utilized a series of properties described below:

#### Secondary structure element preservation (SSEP)

SSEP was calculated as the ensemble average of *S*(**x**), the fraction of native secondary structure in a conformation **x**. Given a native structure **x**_0_, which in our work corresponds to the initial conformation provided to SAM, we considered all its residues in an α-helical (H) or β-strand (E) state, using assignments from the DSSP^76,79^. We define *S* as:

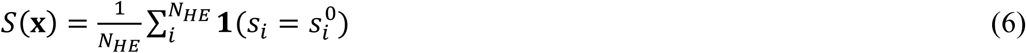

where *N_HE_* is the number of residues in H or E states in **x**_0_, the index *i* is over such residues, *s_i_*^0^ and *s*_*i*_ are the states of residues in **x**_0_ and **x**, respectively, and **1** is an indicator function returning 1 if its equality is satisfied, else 0.

#### Folded state fraction (FSF)

The FSF of an ensemble is based upon *Q*(**x**), the fraction of native contacts^40^ in a conformation. Given a native structure **x**_0_, we considered all contact distances *d*_ij_^0^ ≤ 10 Å between residues *i* and *j* that are at least 3 residues apart in primary sequence. A contact distance is the shortest distance between any heavy atom of *i* and *j*. We used the following definition^26^ for *Q*:

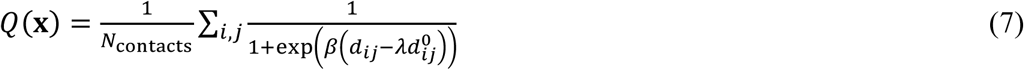

where the *N*_contacts_ is the number of evaluated distances, *d*_*ij*_ is a contact distance in **x**, *β* and *γ* are constants set to 5.0 Å^−1^ and 1.2, respectively. A conformation was considered folded if *Q*(**x**) exceeds a threshold *Q*_thresh_. The FSF of an ensemble is the fraction of conformations with *Q*(**x**) > *Q*_thresh_. While this threshold is usually system-dependent, here we always set it to 0.6. This approximation is justified by two reasons. First, for the purpose of comparing aSAMt and MD ensembles, any reasonable *Q*_thresh_ allows a meaningful analysis of the underlying distributions of *Q*(**x**). Secondly, in mdCATH test MD data, we observed a PCC of 0.92 between FSFs and SSEPs, confirming that for most ensembles 0.6 acceptably expresses foldedness.

#### Melting points

To extract *T̂*_m_ values, we fitted a sigmoid function^42^ to folded state fractions or SSEP values generated by the model. We performed via SciPy^80^ a non-linear least square fitting of:

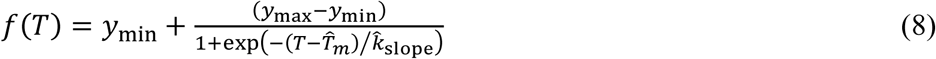

where *y*_min_ and *y*_max_ are the minimum and maximum values in the series generated by aSAMt and *T̂*_*m*_ and *k̂*_slope_ are the parameters to fit.

### Long MD trajectories of fast folding proteins

The sequences of the 12 fast folding proteins were obtained from Lindorff-Larsen et al.^12^. The input 3D conformations for aSAMt originate from the PDB (**Supplementary Table 4**). Wherever possible, we used the raw PDB structures, but for five systems we mutated some residues with MODELLER^81^ to match the original sequences.

The MD trajectories were kindly provided by D.E. Shaw Research. Free energy landscapes were constructed via TICA dimensionality reduction^82^ on MD data, using the PyEMMA library^83^. The input features of TICA were Cα-Cα distances from all pairs of non-adjacent residues. We used system-specific lag times to qualitatively reproduce landscapes from Lewis et al.^26^ (**Supplementary Table 4**). aSAMt ensembles were projected on the axes obtained from TICA on MD data. To compute free energies *G*, we constructed 2D histograms with 50 × 50 uniform bins on the first two TICA axes and applied *G* = −*RT*ln*P*, where *P* are raw histogram frequencies. This is justified by the fact that the long MD trajectories are likely close to equilibrium. To speed up analysis, aSAMt snapshots were not energy-minimized, because TICA was performed only on Cα atoms and the minimization does not change their positions significantly (**Supplementary Fig. 1**).

Data of additional MD trajectories of fast folding proteins^49^ were downloaded from https://github.com/torchmd/torchmd-protein-thermodynamics.

### Proteins with experimentally observed thermal behavior

The experimental data shown in **Fig. 7a** was collected from the S^[Temp]^ dataset in Pucci et al.^52^, containing 222 proteins with *T*_m_ measured through differential scanning calorimetry or circular dichroism. From this set, we selected 62 monomeric proteins meeting specific biomolecular criteria that allowed meaningful comparison with aSAMt (e.g.: no disulfide bridge, cofactors and high sequence similarity with training set proteins). The five pairs of homologous structures with experimental *T*_m_ were collected from Razvi and Schotlz^54^. Details on the selection of these proteins and the setup of their input structures for aSAMt are found in **Supplementary Note 3**.

The nine RFdiffusion thermostable monomers of 200 and 300 residues were originally described in the article of the method^57^ (**Supplementary Fig. 17**). We obtained their AF2 models from: https://figshare.com/s/439fdd59488215753bc3. Control models from the AlphaFold Database^58^ were selected as follows. For each RFdiffusion monomer, we searched in the Swiss-Prot set for proteins of the same length and with pLDDT ≥ 80. We then kept only structures having an absolute difference in *R*_g_ ≤ 0.1 nm with respect to the monomer (i.e.: having the same compactness level). Among these, we selected the structure with the lowest difference in secondary structure composition, evaluated as the L2 distance between two-dimensional vectors with helix and sheet frequencies.

Temperature ranges and other details for the generation of the melting curves of all these proteins are reported in **Supplementary Table 5**.

### Model implementations, software and availability

All neural networks were implemented in PyTorch 1.31.1^84^. Key parts of the autoencoder implementation, such as the Structure Module and *L*_rec_ functions, were adopted from the OpenFold code^72^. Multi-GPU training was performed via PyTorch Lightning. Data analysis was performed with Python scientific libraries^80,85,86^ and MDTraj^76^. Protein structures were rendered with PyMOL.

The code and weights for aSAM models are available at https://github.com/giacomo-janson/sam2. The training, validation and test splits used for mdCATH, as well as input PDB files for the systems that we analyzed are also available in this repository.

## ACKNOWLEDGEMENT

Funding was provided by the National Institutes of Health, grant R35 GM126948 (to MF). We thank D.E. Shaw Research for having kindly shared their fast-folding protein MD data. This work used Bridges-2 (at Pittsburgh Supercomputing Center) and Expanse (at San Diego Supercomputer Center) computing resources through allocation BIO230084 from the Advanced Cyberinfrastructure Coordination Ecosystem: Services & Support (ACCESS) program^87^, which is supported by U.S. National Science Foundation grants #2138259, #2138286, #2138307, #2137603, and #2138296. High-performance computing resources were also used at the Institute for Cyber-Enabled Research (ICER) at Michigan State University.

## AUTHOR CONTRIBUTIONS

GJ, AJ, and MF designed the research, GJ performed and analyzed the machine learning work, AJ performed coarse-grained and atomistic simulations. All authors jointly discussed the findings and wrote the manuscript.

