## Supplementary Figures, Tables, Notes for "Deep generative modeling of temperature-dependent structural ensembles of proteins"

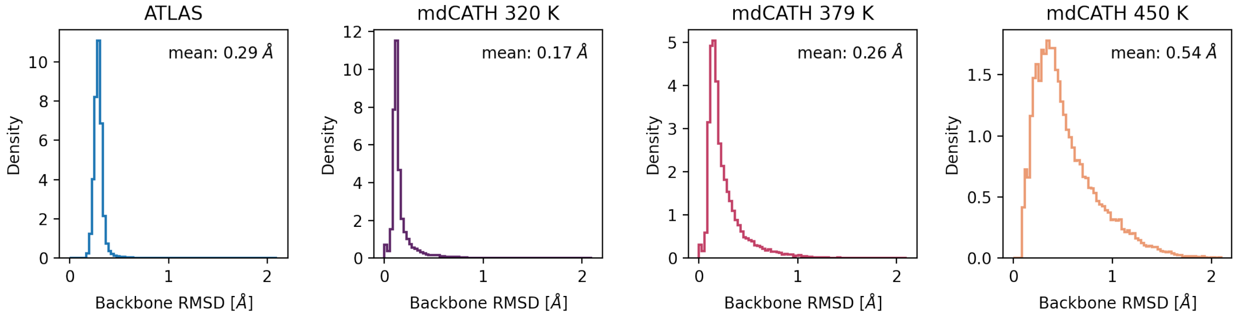


**Supplementary Fig. 1. Impact of the energy minimization process on conformations from aSAM.** For all proteins in the ATLAS and mdCATH test sets, we used of 250 snapshots directly generated by aSAM and applied our energy minimization algorithm on them. We then calculated the backbone RMSD between the original aSAM conformations and their counterparts after minimization. The histograms show the distributions of RMSD values for all snapshots across all test systems. For the ATLAS and mdCATH proteins we used the aSAMc and aSAMt versions described in the main text, respectively.

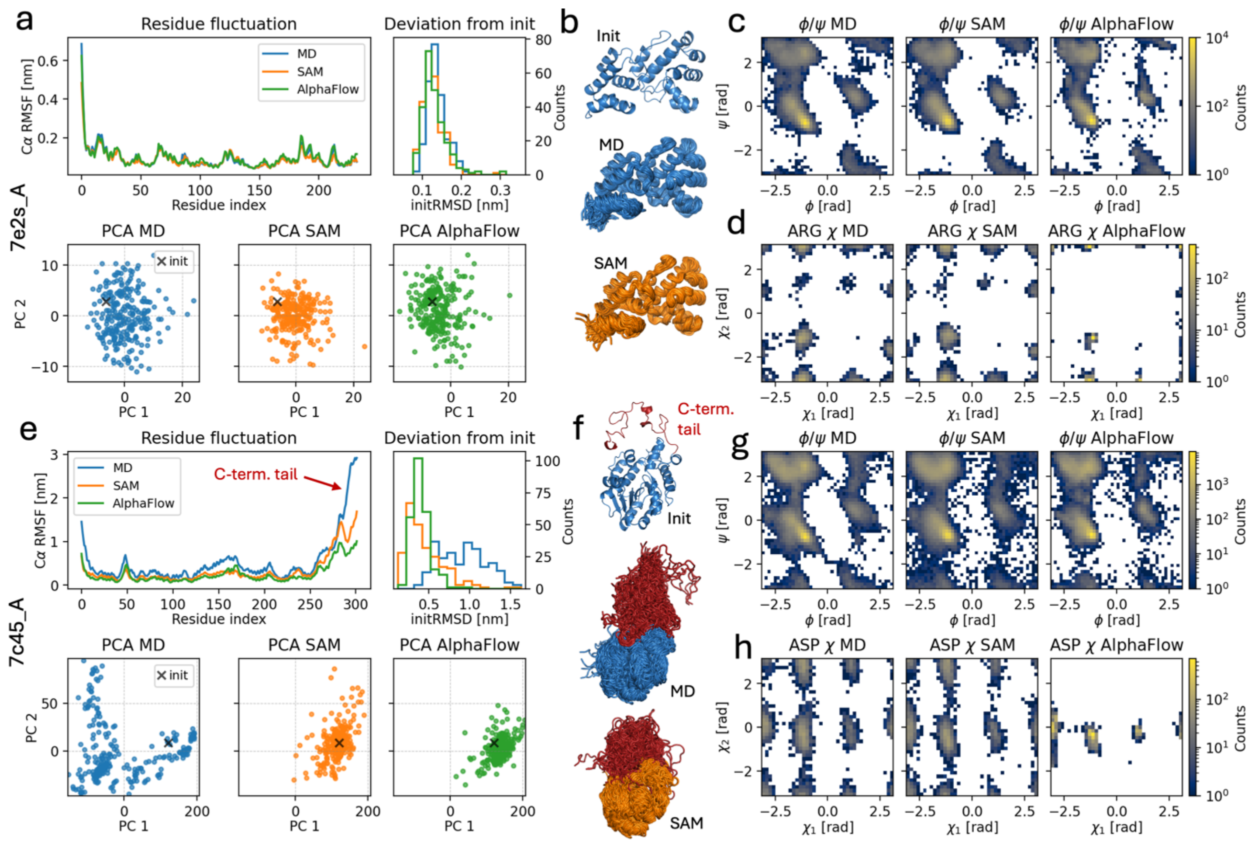


**Supplementary Fig. 2. aSAMc and AlphaFlow ensembles for additional ATLAS test set proteins. a** to **d**: Properties of MD (blue), aSAMc (orange) and AlphaFlow (green) ensembles of test chain 7e2s_A. See **Fig. 2** in the main text for more details. **e** to **h**: Properties of MD and ML ensembles of test chain 7c45_A. **f**: The 7c45_A chain has a C-terminal tail of approximately 50 residues (colored in red). In the MD ensemble, it samples a variety of conformations, while in the ML ensembles it remains close to the initial structure, as seen in initRMSD histograms and PCA, where the two principal components prevalently capture tail dynamics. **d** and **h**: histograms for $\chi_1/\chi_2$ angles for all arginine and aspartate residues in the proteins, respectively. We showcase different amino acids types to demonstrate how aSAMc captures side chain torsions.

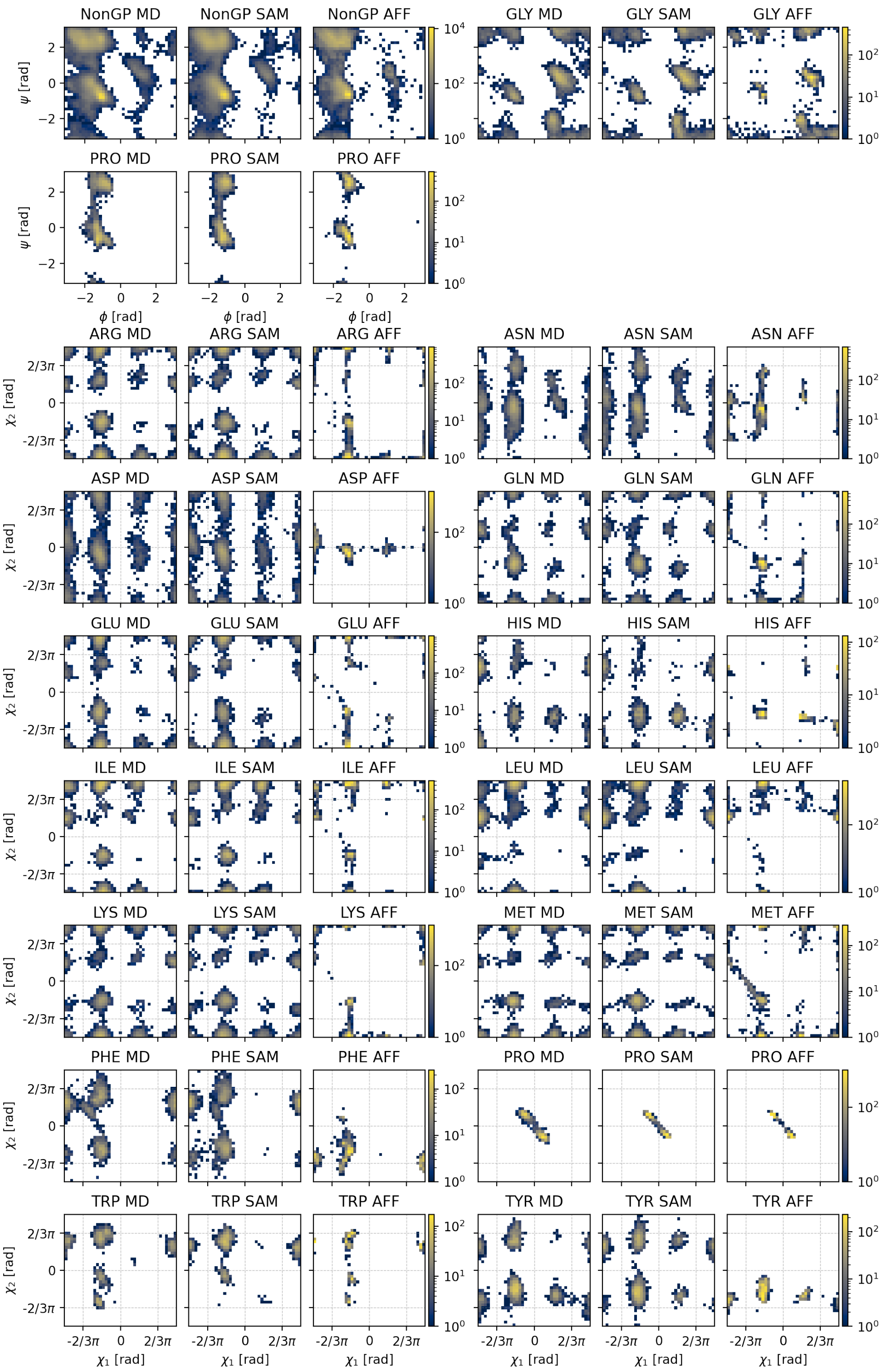


**Supplementary Fig. 3. Torsion angles for the ATLAS test set protein 7p41_D. a** $\phi/\psi$ backbone angles for MD, aSAMc and AlphaFlow (AFF). The histograms report data for all non-proline and non-glycine residues (NonGP), for proline (PRO) and glycine (GLY) residues. **b** Histograms of $\chi_1/\chi_2$ side chain angles for all amino acid types having a pair of those angles. The gray grid represents the bins used to evaluate chiJSD scores (**Methods**). **a** and **b**: the color bars report raw histogram counts from all residues from ensembles with 250 snapshots.

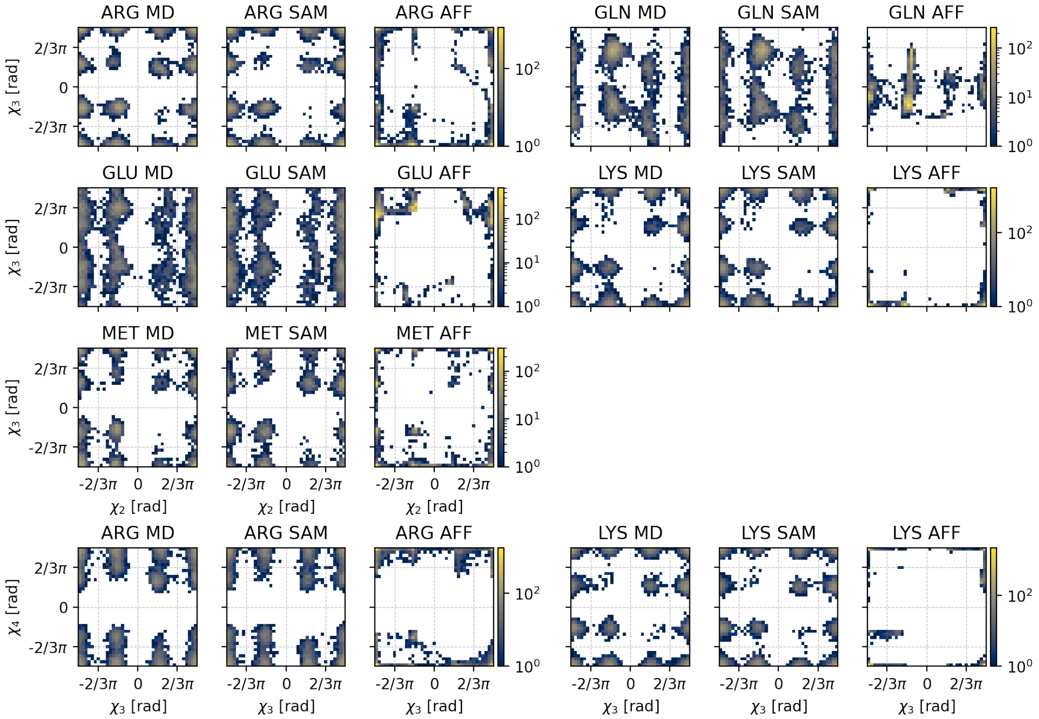


**Supplementary Fig. 4. Additional torsion angle distributions for the ATLAS test set protein 7p41_D.** **a** $\chi_2/\chi_3$ torsion angles for MD, aSAMc and AlphaFlow (AFF). All amino acid types having a pair of those angles are considered. **b** Histograms similarly reporting $\chi_3/\chi_4$ distributions. **a** and **b**: the color bars report raw histogram counts from all sequence positions from ensembles with 250 snapshots. The gray grid represents the bins used to evaluate chiJSD scores.

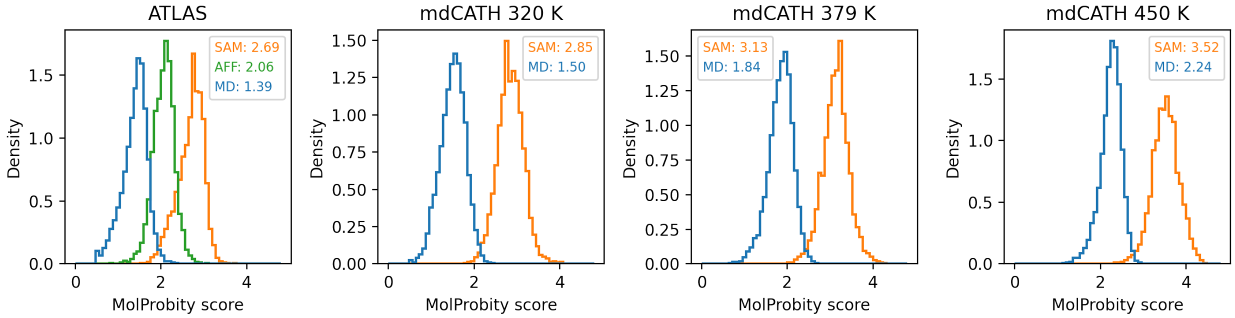


**Supplementary Fig. 5. Distributions of MolProbity scores in MD and ML ensembles.** For all proteins in the ATLAS and mdCATH test sets, we sampled 100 snapshots from their MD (blue), aSAM (orange) and AlphaFlow (AFF, green) ensembles and calculated their MolProbity scores. The histograms show the scores of snapshots from all test ensembles pooled together. Lower scores indicate better stereochemical quality. In the legends, we report the average score of each method beside its name. aSAM scores tend to increase with temperature, but also MD snapshots follow this trend. For the ATLAS and mdCATH proteins we used the aSAMc and aSAMt, respectively. All aSAM ensembles used here were energy minimized, as by default.

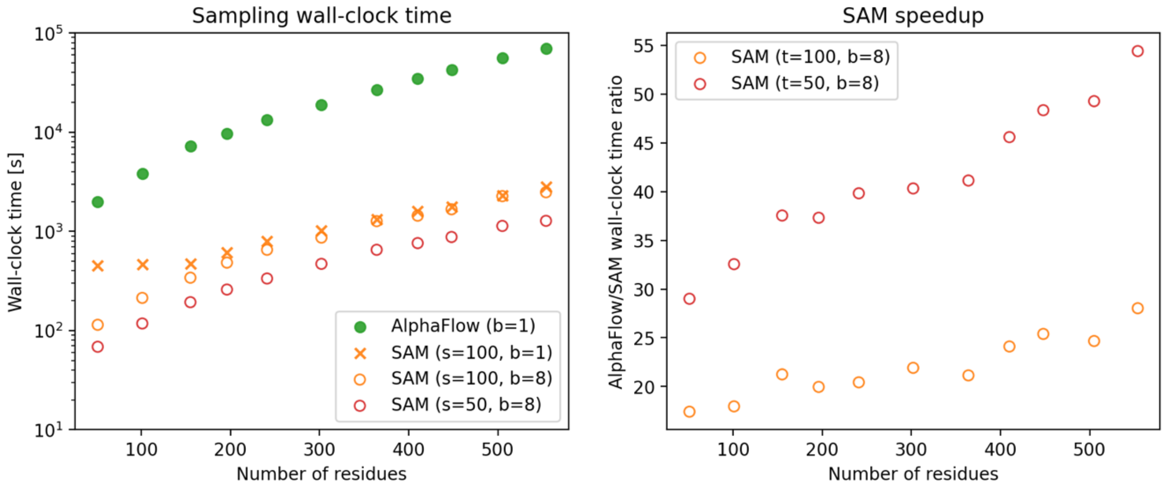


**Supplementary Fig. 6. Wall-clock times of aSAMc and AlphaFlow.** Both ML methods were used to generate ensembles of 250 snapshots for 11 proteins from the ATLAS test set: 6okd_C ($L = 51$), 7n0j_E ($L = 101$), 6q10_A ($L = 155$), 6kty_A ($L = 196$), 6xb3_H ($L = 241$), 7c45_A ($L = 302$), 7aqx_A ($L = 364$), 7onn_A ($L = 410$), 7p41_D ($L = 448$), 7ec1_A ($L = 505$), 6xrx_A ($L = 554$). aSAMc was run with a batch size ($b$ in the legend) of 1 and 8. Note how increasing the batch size in aSAMc leads to efficient parallelization only for small proteins. We used aSAMc with 100 or 50 diffusion steps ($s$ in the legend). The default number of steps in this work is 100. The left panel reports wall-clock times in units of seconds. The right panel reports the speedup of aSAMc with respect to AlphaFlow. All calculations were performed on an NVIDIA V100 GPU. The wall-clock time values are the average measurements from three runs for each protein.

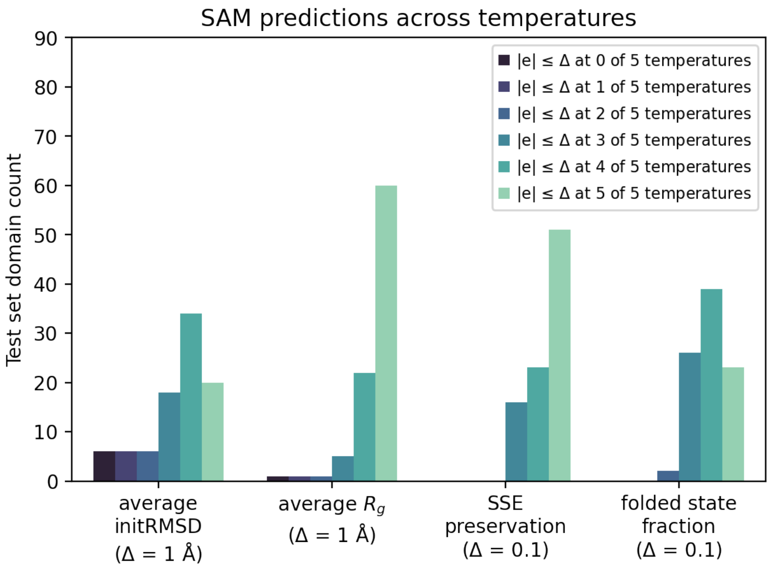


**Supplementary Fig. 7. aSAMt approximations of MD properties of mdCATH across temperatures.** For each property described in **Fig. 3** in the main text, we count the number of test set domains (total number $n = 90$) for which an aSAMt prediction matches an MD value within a threshold Δ across 0, 1, 2, 3, 4 or 5 temperatures.

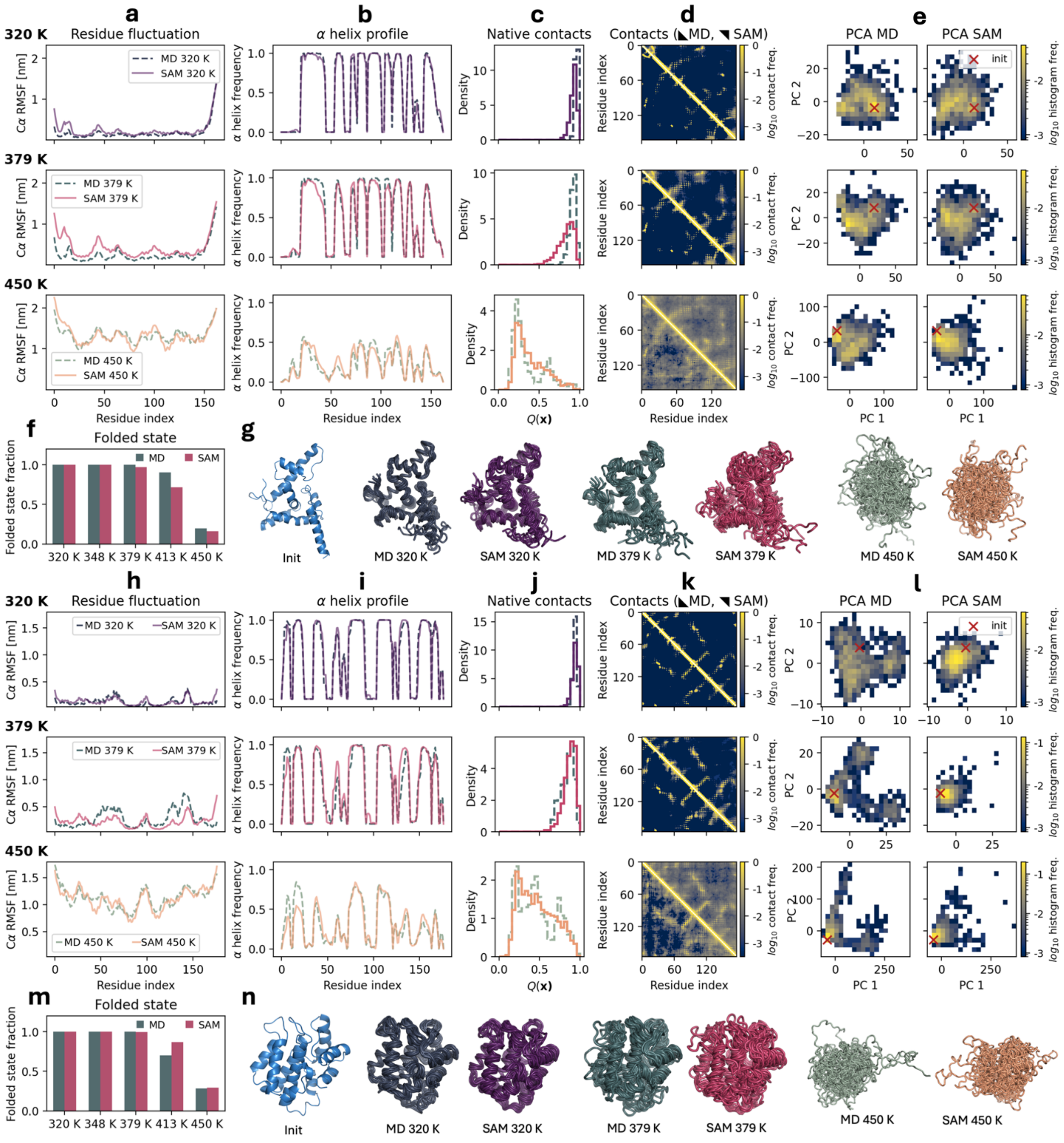


**Supplementary Fig. 8. Properties of the 2vy2A00 and 3nb2A04 test set domains from mdCATH.** These domains are examples of larger more thermostable domains from mdCATH. See **Fig. 4** in the main text for more information. **a-g**: data for 2vy2A00. **h-n**: data for 3nb2A04.

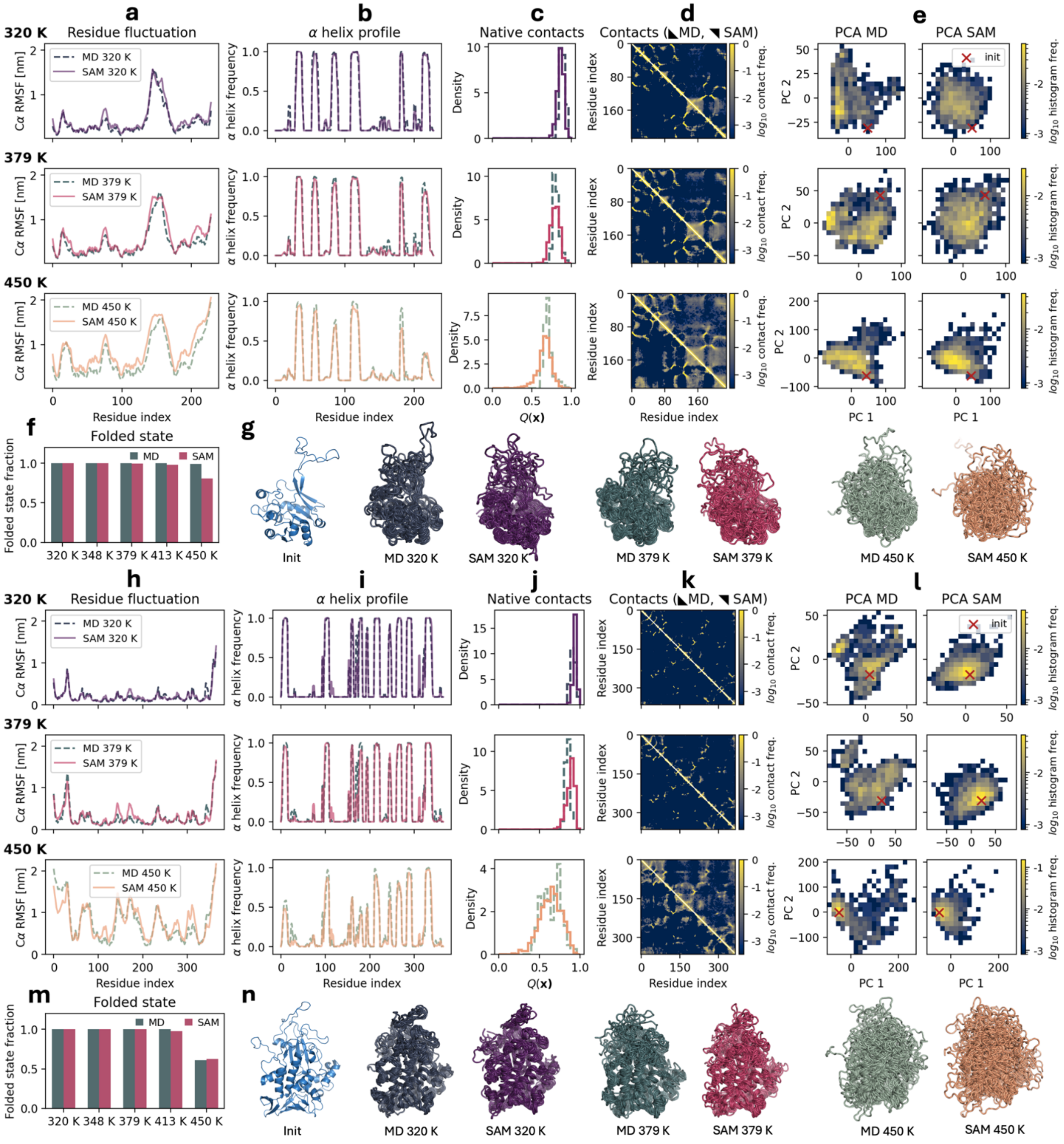


**Supplementary Fig. 9. Properties of the 1qwjB00 and 4a57A02 test set domains from mdCATH.** These domains are examples of larger more thermostable domains from mdCATH. See **Fig. 4** in the main text for more information. **a-g**: data for 1qwjB00. **h-n**: data for 4a57A02.

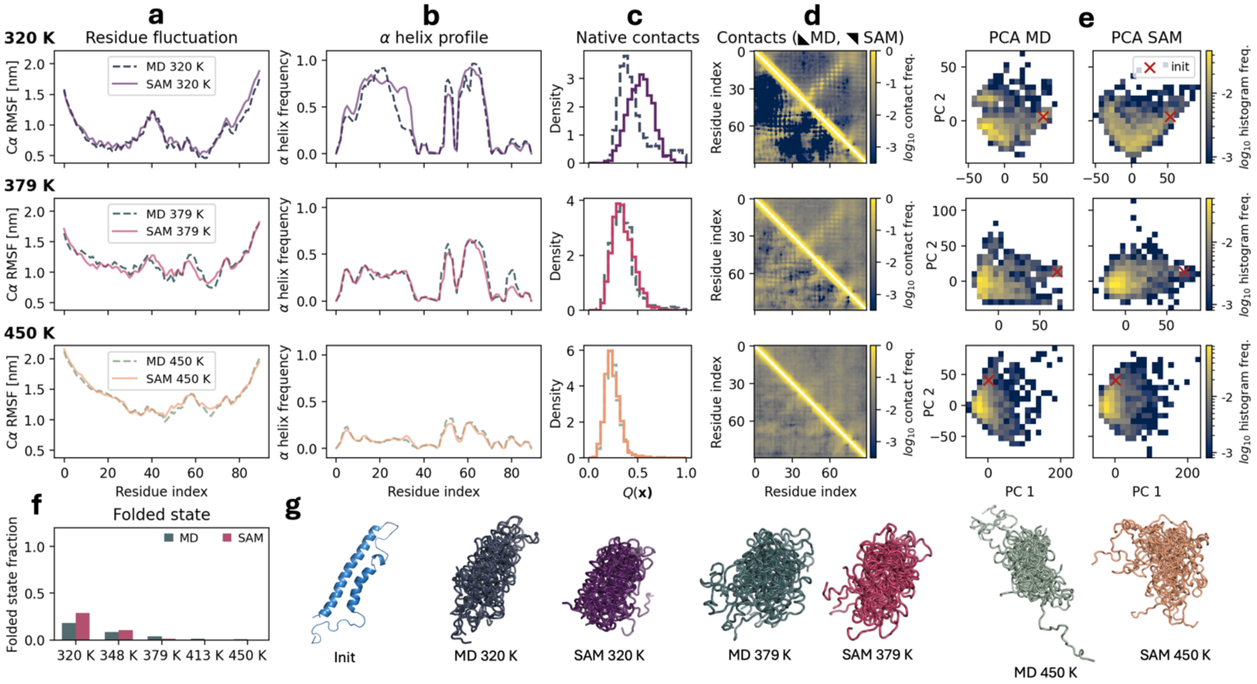


**Supplementary Fig. 10. Properties of the 3nb0A03 test set domain from mdCATH.** This domain is an example of highly dynamical and unstructured domain from mdCATH. See **Fig. 4** in the main text for more information.

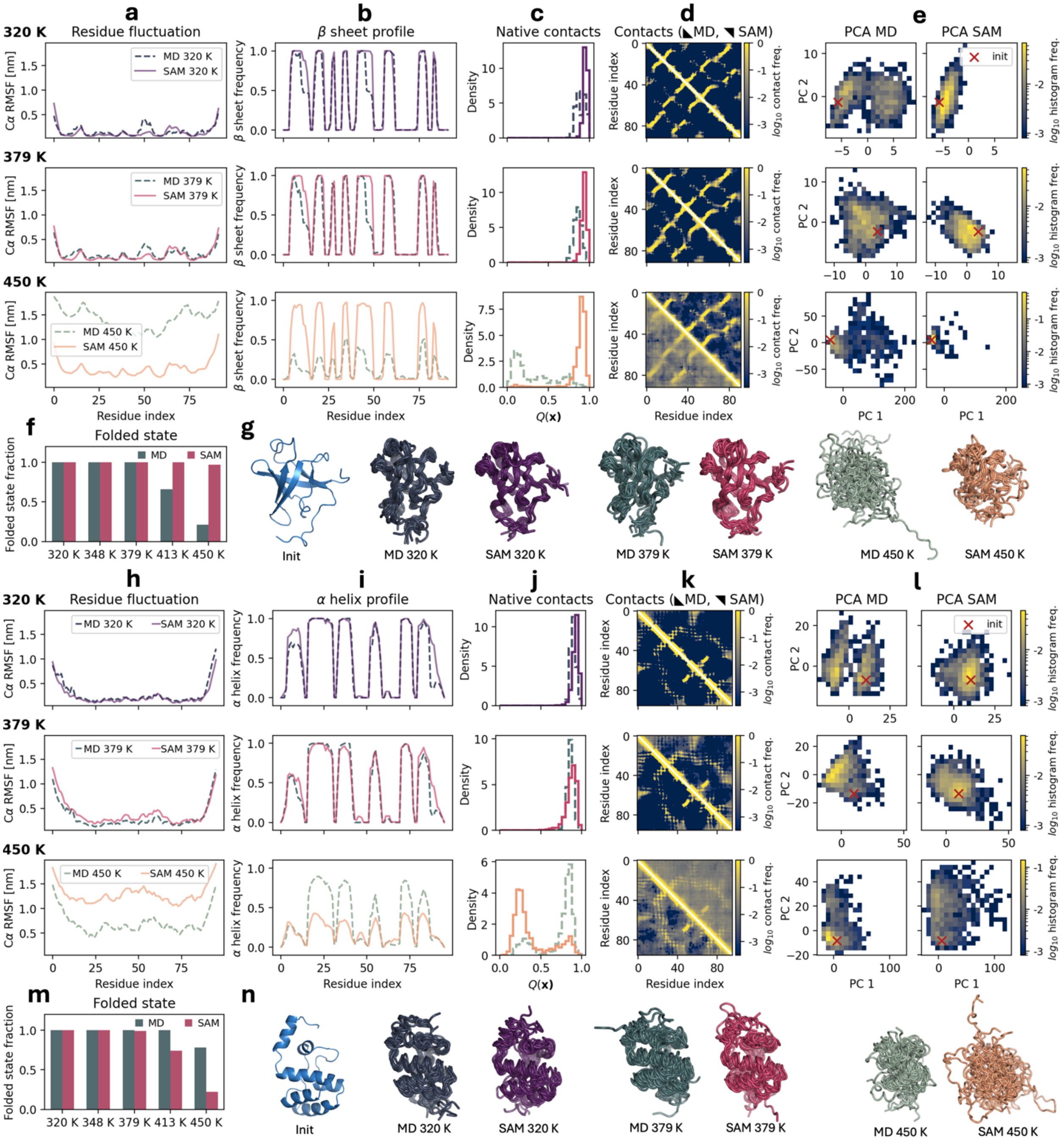


**Supplementary Fig. 11. Properties of the 3u28C00 and 3zrhA01 test set domains from mdCATH.** These two domains are examples of large errors from aSAMt. See **Fig. 4** in the main text for more information. **a-g**: data for 3u28C00. **h-n**: data for 3zrhA01.

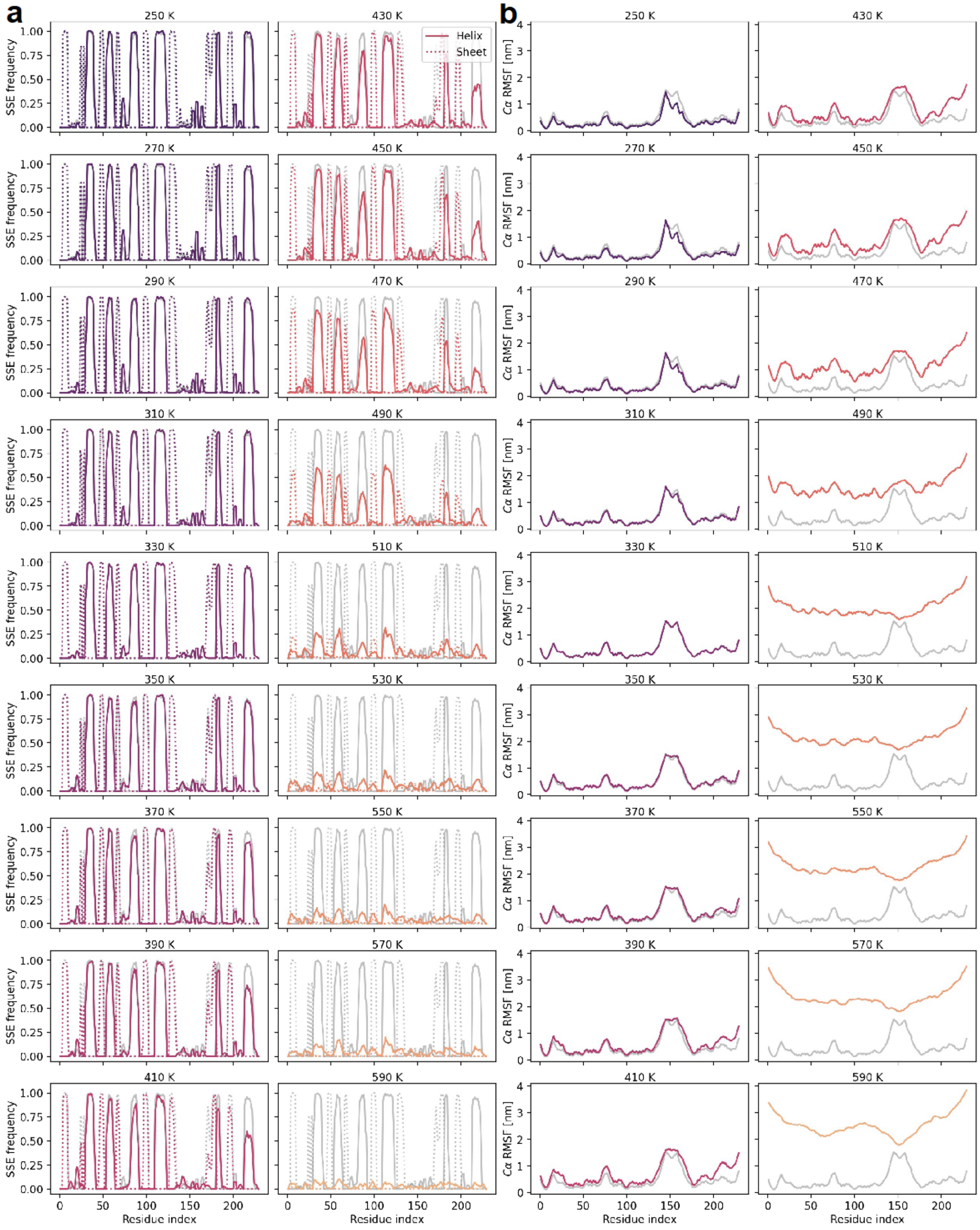


**Supplementary Fig. 12. Properties of aSAMt ensembles for 1qwjB00 across temperatures. a** secondary structure element profiles, with continuous lines for helices and dotted lines for β strands. **b** Cα RMSF profiles. **a-b**: the gray lines represent the profiles at 330 K used for comparing behavior across temperatures.

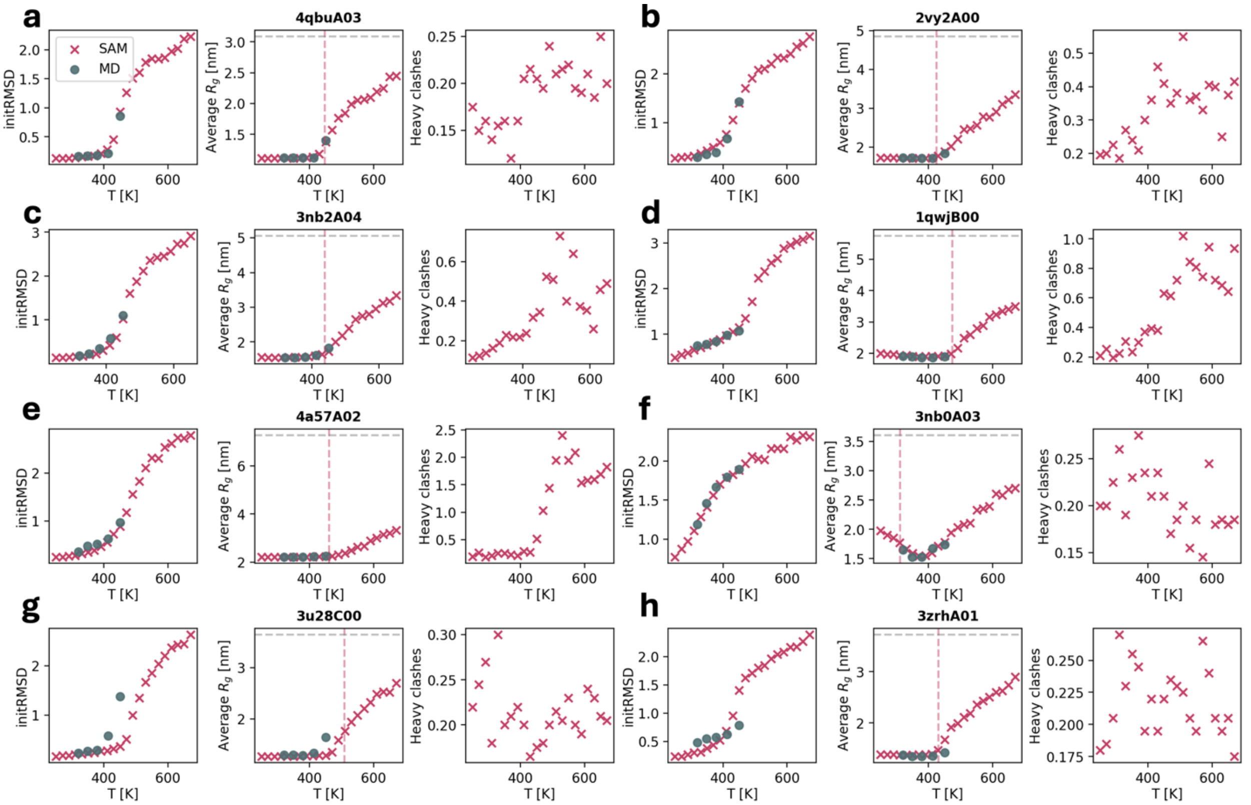


**Supplementary Fig. 13. Additional ensemble properties by scanning across temperatures.** We analyze the same 8 mdCATH test domains shown in **Fig. 5**. In the left subpanels, we show average initRMSD values. In the central subpanels, we show average $R_g$ values. For each domain, the $R_g$ of an ideal non-interacting chain (in $\theta$ solvent) of the same length is shown as a dashed gray horizontal line on the top. The dashed vertical red lines correspond to the $\hat{T}_m$ calculated via FSF values (**Fig. 5**). In the right subpanels, we show the number of heavy clashes per snapshot (**Methods**). aSAMt and MD values are plotted as red and gray markers, respectively. We do not show the number of clashes in MD ensembles, because it is always 0. **a-c** 4qbuA03, 2vy2A00 and 3nb2A04, **d-e** 1qwjB00 and 4a57A02, **f** 3nb0A03, **g-h** 3u28C00 and 3zrhA01.

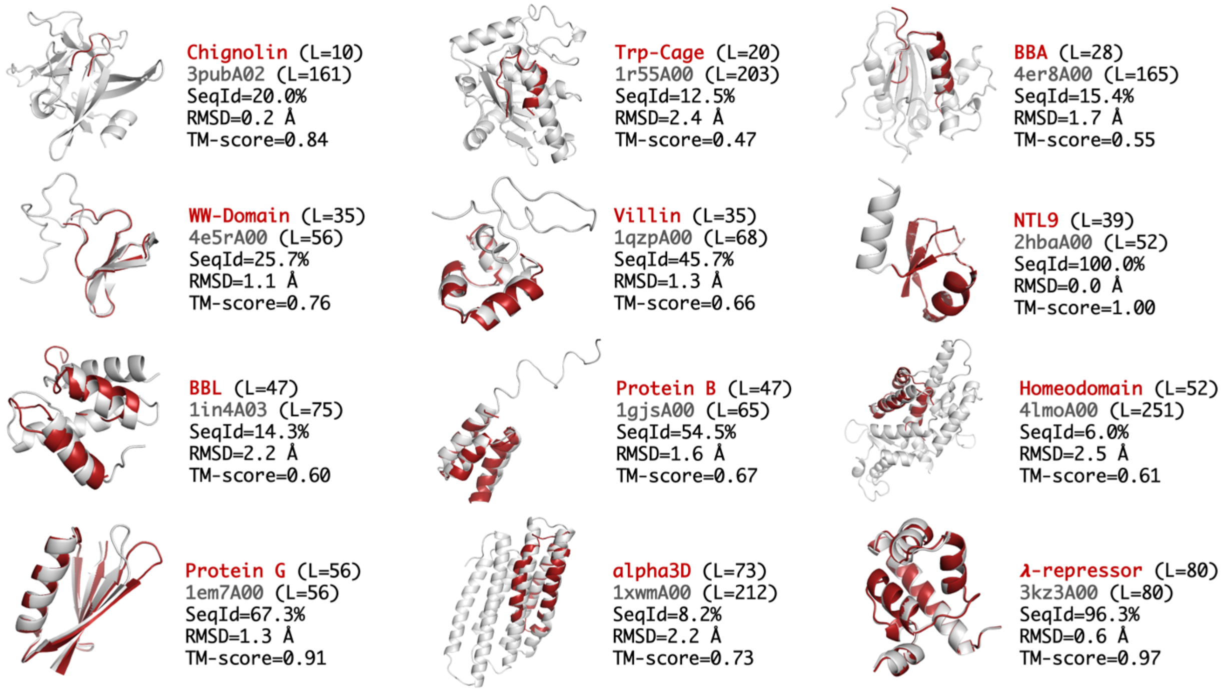


**Supplementary Fig. 14. mdCATH training domains with the highest structural similarity with the fast protein proteins.** For each fast folding protein initial 3D structure (**Supplementary Table 4**), we searched in the aSAMt training set for the most similar domain in terms of TM-score with TM-align[1]. For mdCATH domains, we used the raw CATH structure provided in mdCATH. The initial 3D structures of the fast folders are shown in red, while the most similar domain in the training set is shown in gray. We report the lengths *L* of the two chains, their sequence identity, Cα RMSD and TM-score (normalized by the length of the fast folder chain) in the TM-align alignment.

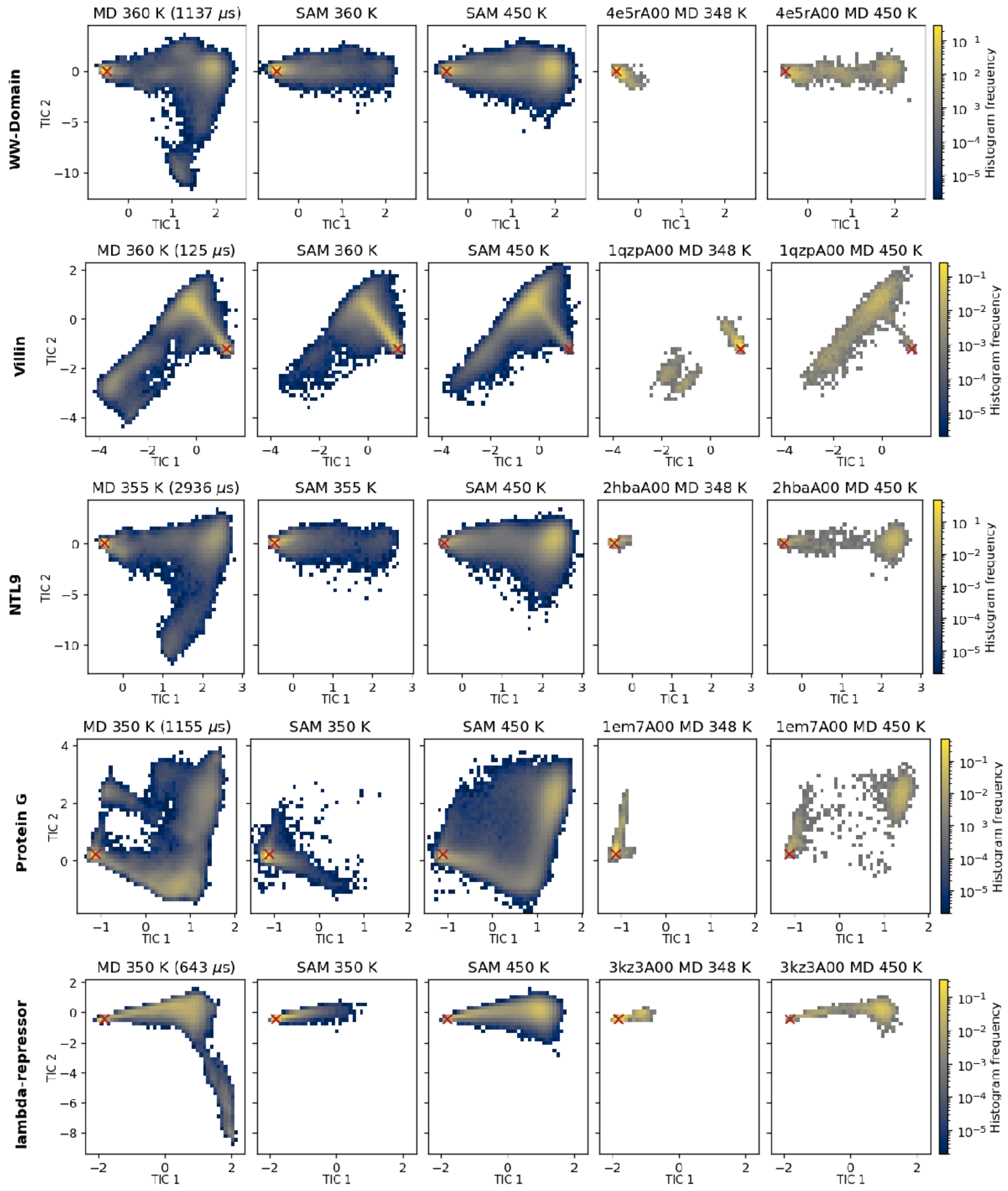


**Supplementary Fig. 15. Comparison from long MD, aSAMt and mdCATH data for 5 fast folding proteins.** Conformations from long MD trajectories, aSAMt and mdCATH ensembles (last two columns) are projected onto TICA axes built from long MD. For each protein, we analyze the mdCATH training domain with the highest structural similarity (with sequence identity ≥ 25% and TM-score ≥ 0.6, **Supplementary Fig. 14**). When a mdCATH domain had different number of residues, we analyze only its aligned residues. Since TICA was based on Cα atoms, mdCATH snapshots remain mappable in presence of residue type differences. Protein B and its closest mdCATH domain were excluded because of gaps in their alignment, which prevented TICA projection. Long MD and aSAMt ensembles have 500k snapshots, the mdCATH ensembles contain all available snapshots in the dataset (≤ 2500). The red × show the location of the input structure of aSAMt. Color bars report the raw histogram frequencies for the data.

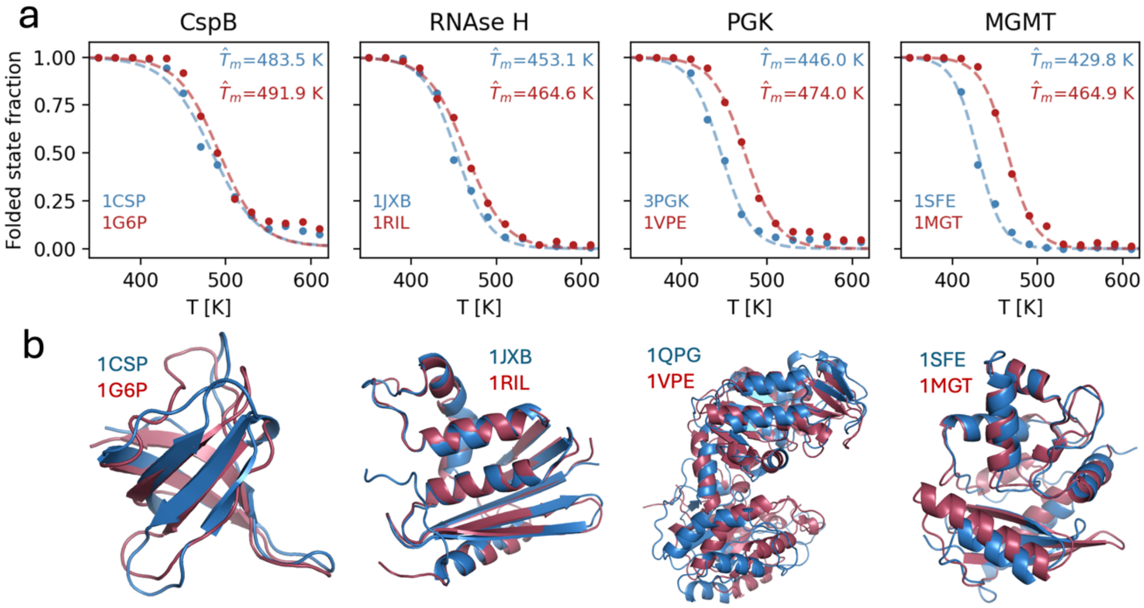


**Supplementary Fig. 16. Additional pairs of homologous proteins with different experimental $T_m$ values. a** aSAMt melting curves for the four pairs of proteins (see **Supplementary Table 6** for more information). Red datapoints correspond to the protein with higher experimental $T_m$ value, blue datapoints to the protein with the lower experimental $T_m$ value. Each plot shows the aSAMt $\hat{T}_m$ values. **b** Structures and PDB ids for the proteins analyzed in **a**.

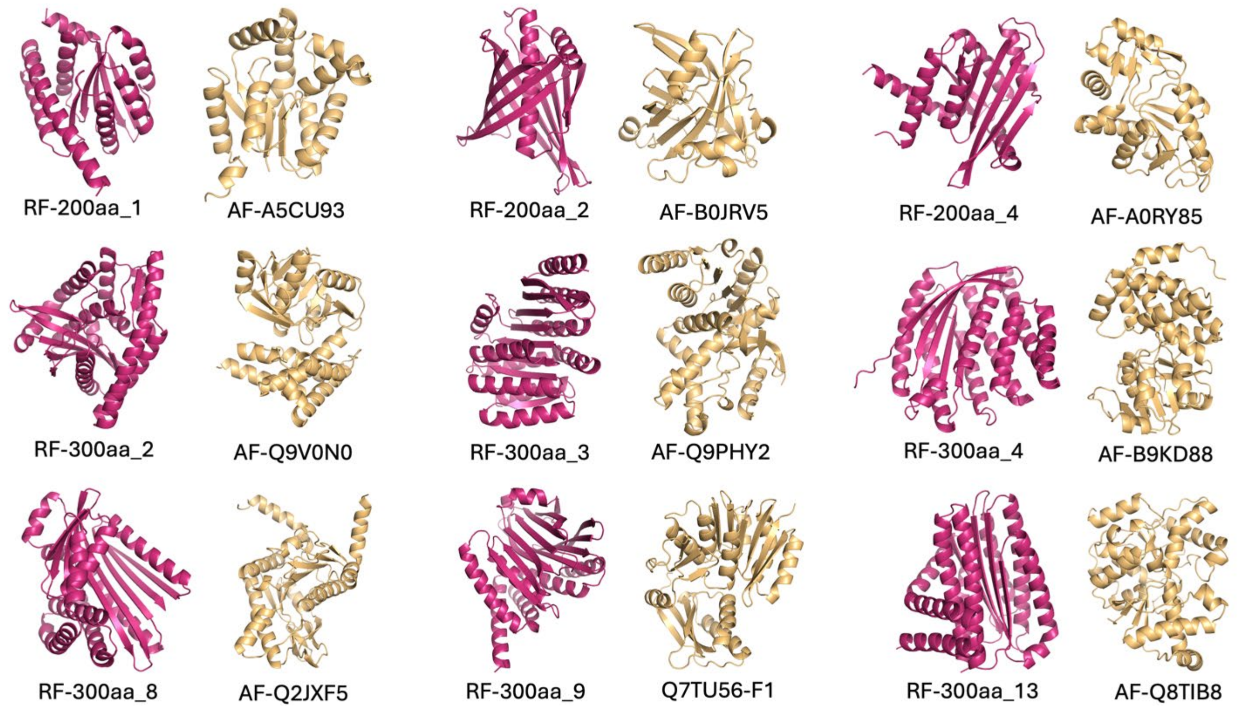


**Supplementary Fig. 17. AF2 models for thermostable RFdiffusion proteins and control proteins from AlphaFold Database.** The 3D model of each RFdiffusion monomer (pink) is shown beside its closest match in the Swiss-Prot dataset (light yellow).

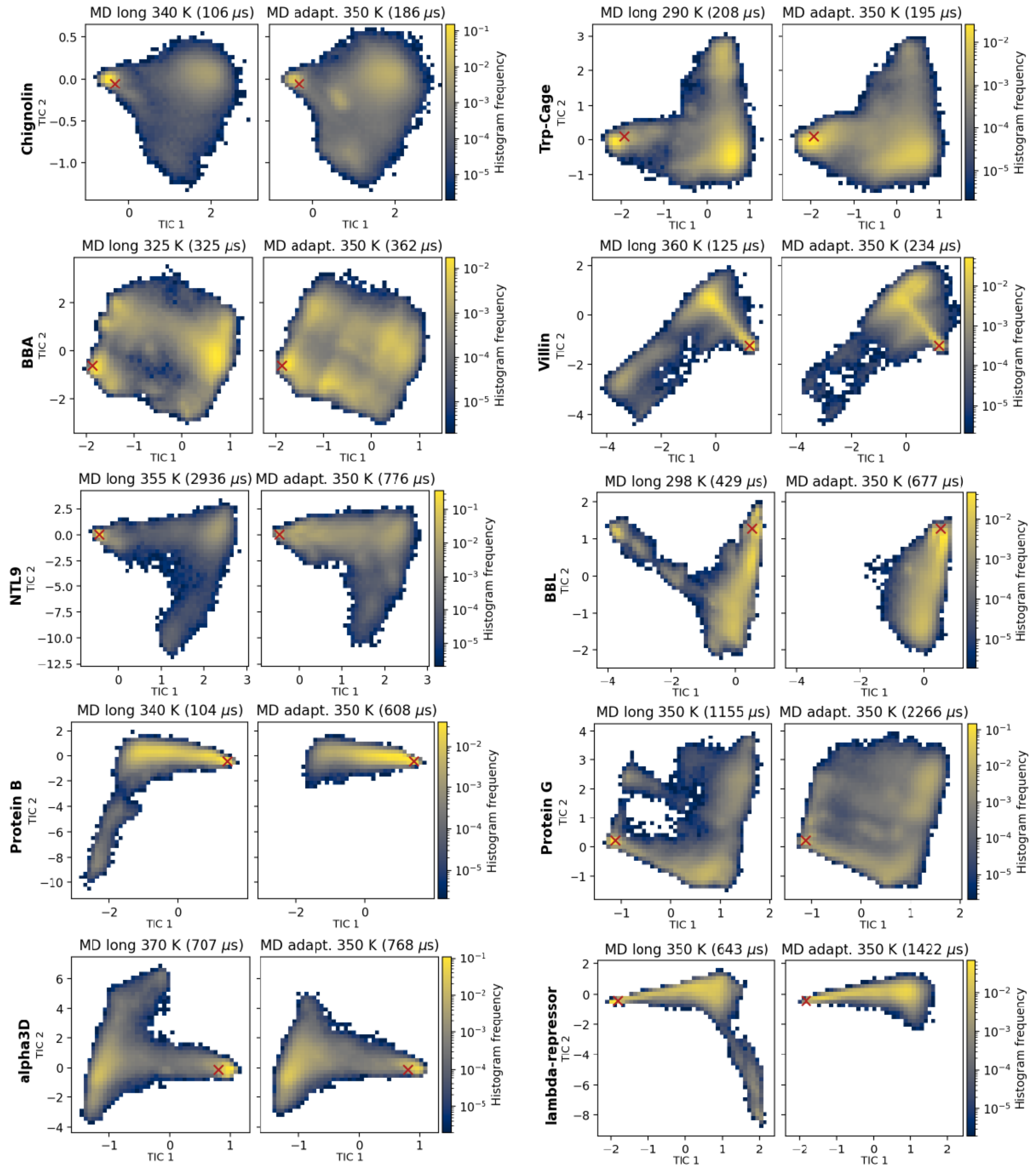


**Supplementary Fig. 18. Comparison of fast folder simulations from a few long MD trajectories and thousands of short ones.** Conformations from adaptive sampling performed in Majewski et al.[2] (right panels) are projected onto TICA axes built upon long simulations by Lindorff-Larsen et al.[3] at D.E. Shaw Research (left panels). Cumulative simulation times reported in the original publications are shown in parentheses. For each dataset, we used 500k randomly extracted snapshots. The red × markers indicate the location of the folded structures of the proteins. Color bars represent raw histogram frequencies. Note that the MD data from Majewski et al. consists of thousands of short trajectories iteratively initiated across the conformational landscapes, leading to density differences compared to long MD due to preferences in the choice of starting points of the short runs. To compare free energies from the two simulation types, the adaptive MD data should be reweighted by Markov state modeling, something we omit here.

**Supplementary Table 1.** Reconstruction ability of the AE models in terms of heavy atom RMSD.

| AE model trained on | ATLAS[a] | mdCATH[b] 320 K | mdCATH[b] 348 K | mdCATH[b] 379 K | mdCATH[b] 413 K | mdCATH[b] 450 K |
|---|---|---|---|---|---|---|
| ATLAS | 0.38 ± 0.02 | 0.28 ± 0.01 | 0.29 ± 0.01 | 0.30 ± 0.005 | 0.32 ± 0.004 | 0.40 ± 0.02 |
| mdCATH 320-450 K | 0.33 ± 0.01 | 0.29 ± 0.01 | 0.30 ± 0.01 | 0.31 ± 0.01 | 0.34 ± 0.004 | 0.42 ± 0.02 |

The AE models trained on ATLAS and mdCATH were evaluated on ATLAS test set chains and mdCATH test set domains at five different temperatures. The table shows heavy atom RMSD values (in Å) between MD snapshots and their autoencoded versions. The latter were obtained by encoding MD snapshots with the encoder and reconstructing them with the decoder. For each test system, ensembles of 250 snapshots were randomly extracted from MD data and the ensemble average reconstruction RMSD was computed. The table reports the mean, along with standard errors, of these ensemble averages across all test systems.

[a]Evaluation on ATLAS test set proteins ($n$ = 82).
[b]Evaluation on mdCATH test set domains ($n$ = 90).

**Supplementary Table 2.** Effect of the number of reverse diffusion steps in aSAMc, evaluated on ATLAS MD ensembles.

| **Strategy** | **PCC Cα RMSF (↑)** | **WASCO-glob (↓)** | **WASCO-loc (↓)** | **chiJSD (↓)** | **Heavy clashes (↓)** | **Peptide bond length violations (↓)** |
|---|---|---|---|---|---|---|
| $s$=100 | 0.886 ± 0.011 | 158.2 ± 16.8 | 1.15 ± 0.05 | 0.07 ± 0.002 | 0.23 ± 0.04 | 0.52 ± 0.09 |
| $s$=50 | 0.887 ± 0.011 | 159.1 ± 16.9 | 1.15 ± 0.05 | 0.07 ± 0.002 | 0.25 ± 0.05 | 0.54 ± 0.09 |
| $s$=40 | 0.888 ± 0.010 | 159.8 ± 17.0 | 1.15 ± 0.05 | 0.07 ± 0.002 | 0.24 ± 0.04 | 0.52 ± 0.09 |
| $s$=25 | 0.885 ± 0.011 | 161.1 ± 17.0 | 1.16 ± 0.05 | 0.07 ± 0.002 | 0.23 ± 0.04 | 0.53 ± 0.09 |
| $s$=10 | 0.875 ± 0.011 | 165.6 ± 17.5 | 1.19 ± 0.05 | 0.08 ± 0.002 | 0.32 ± 0.07 | 0.57 ± 0.09 |
| $s$=5 | 0.805 ± 0.019 | 174.1 ± 20.8 | 1.22 ± 0.05 | 0.08 ± 0.002 | 1.62 ± 0.44 | 0.93 ± 0.15 |

In the **Strategy** column, $s$ is the number of reverse diffusion steps when sampling with aSAMc. See **Table 1** in the main text for more information on the dataset and the meaning of columns. The first two rows here are replicated from that table for convenience.

**Supplementary Table 3.** GPU wall-clock times for MD simulations and sampling via aSAMt.

| Domain | MD simulation efficiency [ns/day] | MD wall-clock time for 2.5 μs of simulation [s] | aSAMt wall-clock time for generating 500 snapshots [s] |
|---|---|---|---|
| 4qbuA03 (*L*=66) | 227.0 | $9.5 \times 10^5$ | $4.1\times 10^2$ |
| 4a57A02 (*L*=366) | 63.3 | $3.4 \times 10^6$ | $3.4 \times 10^3$ |

Two mdCATH test set domains were analyzed. Their names and number of residues *L* are shown in the **Domain** column. Explicit solvent MD was carried out with a protocol comparable to the one used in mdCATH. Simulations were performed in OpenMM 8.0.0 using the CHARMM36m force field[4] with a 2 fs integration time step in a box extending 15 Å beyond the solute. Particle Mesh Ewald was applied with a 12 Å cutoff. van der Waals interactions were switched off between 10 and 12 Å. Simulations were GPU-accelerated on an NVIDIA RTX 2080 Ti. The systems contained 19,777 atoms (4qbuA03) and 83,414 atoms (4a57A02). We measured the simulation efficiency (ns/day) in limited-length trajectories and extrapolated the time required to obtain 2.5 μs of simulation time. The aSAMt ensembles were generated on a RTX 2080 Ti GPU, with batch sizes of 4 for sampling and 50 for energy minimization, respectively. The aSAMt wall-clock times are the average from three runs for each domain.

**Supplementary Table 4.** Description of the proteins, MD data and its analysis for the 12 fast folder simulations from D.E. Shaw Research.

| Protein | Num. of residues | Initial structure PDB | Num. of MD trajectories | MD $T$ [K] | Total simulation time [µs] | TICA lag time [ns] |
|---|---|---|---|---|---|---|
| Chignolin | 10 | 5AWL | 1 | 340 | 106 | 20 |
| Trp-Cage | 20 | 2JOF* | 1 | 290 | 208 | 20 |
| BBA | 28 | 1FME | 2 | 325 | 325 | 20 |
| WW-Domain | 35 | 2F21* | 2 | 360 | 1137 | 60 |
| Villin | 35 | 2F4K* | 1 | 360 | 125 | 20 |
| NTL9 | 39 | 2HBA | 4 | 355 | 2936 | 20 |
| BBL | 47 | 2WXC | 1 | 298 | 429 | 20 |
| Protein B | 47 | 1PRB* | 1 | 340 | 104 | 40 |
| Homeodomain | 52 | 2P6J | 2 | 360 | 327 | 20 |
| Protein G | 56 | 1MI0 | 4 | 350 | 1155 | 20 |
| alpha3D | 73 | 2A3D | 2 | 370 | 707 | 12 |
| λ-repressor | 80 | 1LMB* | 4 | 350 | 643 | 20 |

*Mutations were introduced in the PDB structure using MODELLER to obtain the same sequence used in Lindorff-Larsen et al.[3]

**Supplementary Table 5.** Parameters used to generate melting curves in Fig. 7 and Supplementary Fig. 17.

| **Dataset** | **Reported in** | **Temperature range [K]** | **Temperature interval [K]** | **Num. of generated conformations per *T*** |
|---|---|---|---|---|
| 62 monomers | Fig. 7a | 250-710 | 20 | 200 |
| 5 homologous pairs | Fig. 7b and Supplementary Fig. 17 | 250-710 | 20 | 200 |
| 9 RFdiffusion monomers and AF2 controls | Fig. 7c | 270-710 | 20 | 250 (5 subsamples of 50) |

**Supplementary Table 6.** Details of pairs of homologous proteins with different experimental $T_m$.

| Group | SeqId[a] % | PDB codes | Organism | SAM $\Delta\hat{T}_m$ [K] | Exp. $\Delta T_m$ [K] |
|---|---|---|---|---|---|
| CspB | 62.1 | 1CSP<br>1G6P | *Bacillus subtilus*<br>*Thermotoga maritima* | 8.4 | 29.3 |
| RNAse[b] H | 54.0 | 1JXB<br>1RIL | *Escherichia coli*<br>*Thermus thermophilus* | 11.5 | 20.0 |
| PGK[c] | 38.4 | 3PGK<br>1VPE | *Saccharomyces cerevisiae*<br>*Thermotoga maritima* | 28.0 | 28.8 |
| RP[d] L30E | 32.3 | 1CN7<br>1H7M | *Saccharomyces cerevisiae*<br>*Thermococcus celer* | 32.6 | 48.1 |
| MGMT[e] | 25.7 | 1SFE<br>1MGT | *Escherichia coli*<br>*Thermococcus kodakiensis* | 35.1 | 54.8 |

For each group, the differences in aSAMt and experimental (Exp.) melting points are calculated between the values of the protein in the second row (red text, higher experimental $T_m$) and the first row (blue text, lower experimental $T_m$). The experimental $T_m$ values were obtained from Razvi & Scholtz 2006[5], which provides references for the articles that originally reported the $T_m$ measurements.

[a]Sequence identity in the TM-align alignment of the two 3D structures.
[b]Ribonuclease (RNAse).
[c]Phosphoglycerate kinase (PGK).
[d]Ribosomal protein (RP).
[e]$O^6$Methyl guanine-DNA methyl transferase (MGMT).

### Supplementary Note 1

### Details of neural networks and their training

#### 1 - Neural networks overview

**aSAM versions**. For the AE models, we used the same hyperparameters for versions trained on ATLAS and mdCATH. The noise prediction networks have minor differences, which will be highlighted below.

**Activation functions**. Unless otherwise stated, all activation functions in aSAM are SiLU.

**Multilayer perceptrons (MLP)**. Unless otherwise stated, all MLPs have the following structure:

$$\mathrm{Linear}(\mathbf{x}, in = \mathrm{dim\_1}, out = \mathrm{dim\_2}) \rightarrow \mathrm{SiLU}(\mathbf{x}) \rightarrow \mathrm{Linear}(\mathbf{x}, in = \mathrm{dim\_2}, out = \mathrm{dim\_2}).$$

**RBFs**. We make use of radial basis function (RBF) expansions to embed inter-atomic distances. We always use the expnorm expansion from TorchMD-NET[6], with a cutoff minimum of 0 nm and varying maxima. We use different numbers of RBF centers and always set their parameters as trainable. To refer to an expansion, we denote it as $\mathrm{RBF}(cutoff\_max, num\_rbfs)$, with the maximum expressed in units of nm.

**Positional embeddings**. In all networks, we make use of learnable relative 2D positional embeddings introduced in AlphaFold2[7], with a maximum sequence separation of 32.

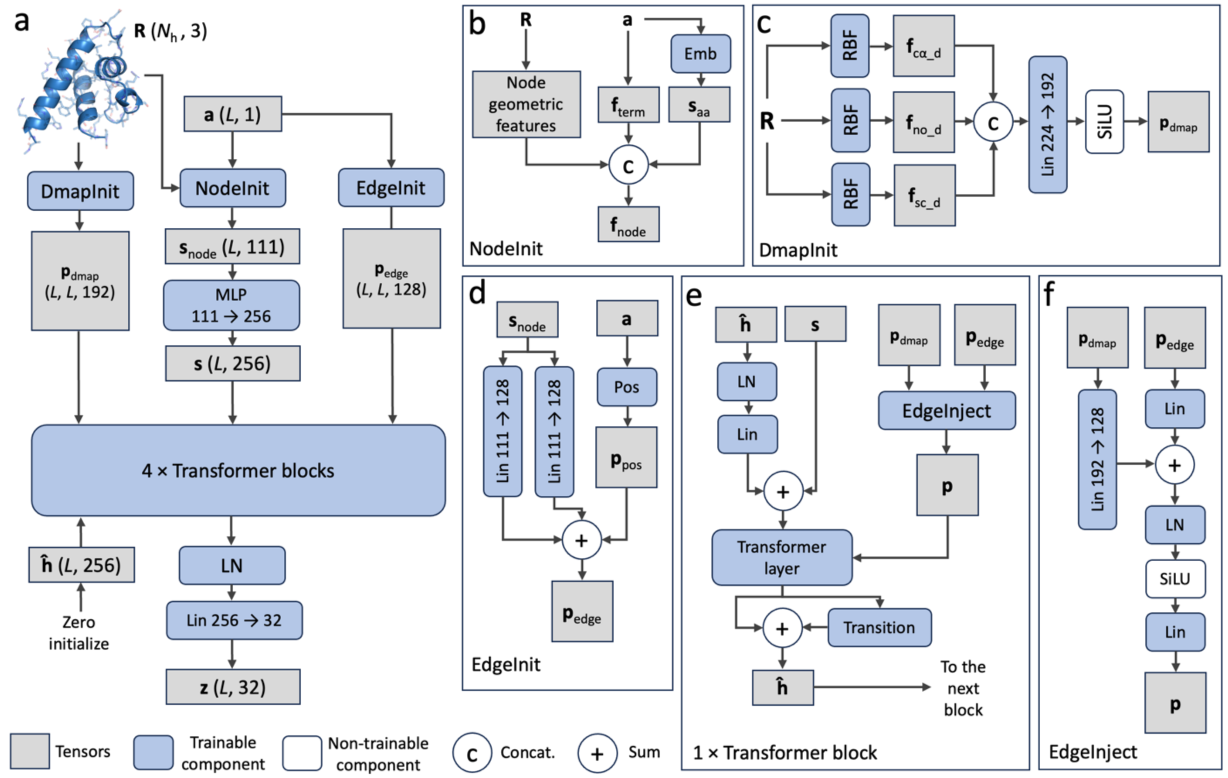


**Supplementary Fig. 19. Architecture of the encoder in aSAM. a** Outline of the encoder. **b-f** Architecture of the neural network modules in the encoder. We use the following abbreviations. Lin *in_dim* → *out_dim*: linear layers with their input and output dimensions (when no dimension value is shown, *in_dim* = *out_dim*, with *in_dim* that can be inferred from previous layers); LN: layer normalization; RBF: radial basis function; Emb: simple token embedding layer; Pos: AlphaFold2-like positional embedding layer; MLP: multilayer perceptrons.

### 2 - Encoder network

#### 2.1 - Overview

**Outline**. The encoder $E_\phi$ has a minorly modified transformer architecture (**Supplementary Fig. 20a**). This architecture was found to improve AE reconstruction in early trials, and we decided to keep it.

**Input**. The input of $E_\phi$ is:

- $\mathbf{R} \in \mathbb{R}^{N_\mathrm{h} \times 3}$: the heavy atom coordinates of a protein chain of length $L$ with $N_\mathrm{a}$ heavy atoms.
- $\mathbf{a} \in \mathbb{R}^{L \times 1}$: the amino acid sequence.

Such inputs are converted into several features used as conditions to update a node representation $\hat{\mathbf{h}} \in \mathbb{R}^{L \times 256}$ initialized with zeros.

#### 2.2 - Node features

**Node geometric features**. The coordinates **R** are converted into roto-translationally invariant node features describing the local geometry of a chain:

- $\mathbf{f}_{\mathrm{c\alpha\_tor}} \in \mathbb{R}^{L \times 3}$: torsion angles formed by four consecutive Cα atoms. The first two channels contain the cosine and sine values of the $L - 3$ angles of a chain. The third channel contains 0 for positions with missing angle values (for residues at the chain termini not having enough neighbors) and 1 for the rest. The channel for labeling missing values is also used in the $\mathbf{f}_{\mathrm{c\alpha\_ba}}$, $\mathbf{f}_{\mathrm{c\alpha\_bl}}$, $\mathbf{f}_\varphi$, $\mathbf{f}_\psi$ and $\mathbf{f}_\omega$ features below.
- $\mathbf{f}_{\mathrm{c\alpha\_ba}} \in \mathbb{R}^{L \times 3}$: bond angles formed by three consecutive Cα atoms. Its first two channels contain the cosine and sine values of the $L - 2$ angles of a chain.
- $\mathbf{f}_{\mathrm{c\alpha\_bl}} \in \mathbb{R}^{L \times 2}$: distances between pairs of adjacent Cα atoms. Its first channel contains the raw values (in units of nm) of the $L - 1$ distances in a chain.
- $\mathbf{f}_\varphi \in \mathbb{R}^{L \times 3}$, $\mathbf{f}_\psi \in \mathbb{R}^{L \times 3}$ and $\mathbf{f}_\omega \in \mathbb{R}^{L \times 3}$: $\varphi$, $\psi$ and $\omega$ backbone torsion angles. Their first two channels contain the cosine and sine values of the $L - 1$ angles of a chain.
- $\mathbf{f}_\chi \in \mathbb{R}^{L \times 12}$: $\chi_1$, $\chi_2$, $\chi_3$ and $\chi_4$ side chain torsion angles. The cosine and sine values of all the 4 possible $\chi$ angles at each residue are stored in 8 channels. Another 4 channels represent missing $\chi$ values with 0 and existing ones with 1.
- $\mathbf{f}_{\mathrm{local\_3d}} \in \mathbb{R}^{L \times 49}$: local atomic coordinates. For each residue $i$, a window of 3 N- and C-terminal residues is taken. The coordinates of the Cα atoms and side chain centroids (SCC) of this window of 7 residues (including $i$) are transformed into the backbone frame of $i$, defined as in AF2. This yields $7 \times 2 \times 3 = 42$ channels. For residues near chain termini, coordinates of missing neighbors are filled with 0. Another 7 channels are included, storing 0 for missing neighbors and 1 for existing ones, leading to 49 channels.

**Node amino acid features**. The amino acid sequence **a** is used to build these features:

- $\mathbf{s}_{\mathrm{aa}} \in \mathbb{R}^{L \times 32}$: mapped from **a** through an embedding layer with learnable parameters.
- $\mathbf{f}_{\mathrm{term}} \in \mathbb{R}^{L \times 1}$: additional feature to label the N- and C-terminal residues of a chain. All positions store 0 and the first and last positions store 1.

**Node embeddings**. All node features from above are concatenated to a node feature tensor $\mathbf{s}_{\mathrm{node}} \in \mathbb{R}^{L \times 111}$ (**Supplementary Fig. 20b**). This is then projected via a MLP to a node embedding $\mathbf{s} \in \mathbb{R}^{L \times 256}$, one of the inputs of the transformer blocks.

##### 2.3 - Edge features

**Edge geometric features**. The coordinates **R** are also converted into distance matrices, used as edge features. These important features describe the global geometry of a protein chain:

- $\mathbf{f}_{\text{c}\alpha\_\text{d}} \in \mathbb{R}^{L\times L\times 128}$: an RBF(10.0, 128) of all Cα-Cα distances.
- $\mathbf{f}_{\text{no\_d}} \in \mathbb{R}^{L\times L\times 32}$: an RBF(3.0, 32) of N-O distances.
- $\mathbf{f}_{\text{sc\_d}} \in \mathbb{R}^{L\times L\times 64}$: an RBF(7.0, 64) of distances between pairs of SCCs.

**Edge amino acid features**. The sequence is used to derive positional embeddings $\mathbf{p}_{\text{pos}} \in \mathbb{R}^{L\times L\times 128}$.

**Edge embeddings**. The RBF expansions are then concatenated and projected to a representation $\mathbf{p}_{\text{dmap}} \in \mathbb{R}^{L\times L\times 192}$ in the DmapInit module (**Supplementary Fig. 20c**). Additionally, node features $\mathbf{s}_{\text{node}}$ and positional embeddings $\mathbf{p}_{\text{pos}}$ are processed by the EdgeInit module to obtain an edge embedding $\mathbf{p}_{\text{edge}} \in \mathbb{R}^{L\times L\times 128}$ (**Supplementary Fig. 20d**).

##### 2.4 - Transformer blocks

**Node representation updates**. The embedding $\mathbf{s}$ is utilized to update the node representation $\hat{\mathbf{h}}$ in a series of 4 transformer blocks (**Supplementary Fig. 20e**). The blocks have a standard transformer multihead self-attention module with 16 heads and a "pre" layer normalization configuration. At each block, $\mathbf{p}_{\text{dmap}}$ and $\mathbf{p}_{\text{edge}}$ are combined in the EdgeInject module and the resulting representation $\mathbf{p} \in \mathbb{R}^{L\times L\times 128}$ is linearly projected to attention biases for updating $\hat{\mathbf{h}}$ (**Supplementary Fig. 20f**). Before being passed at the next block, $\hat{\mathbf{h}}$ is processed via a residual connection and a Transition module from AlphaFold3[8].

**Output**. After the last block, $\hat{\mathbf{h}}$ is mapped via an output module to a SE(3)-invariant encoding $\mathbf{z} \in \mathbb{R}^{L\times 32}$.

#### 3 - Decoder network

The decoder $D_\psi$ is based on a Structure Module from AlphaFold2, with few additional layers for processing its input. The input of $D_\psi$ is an encoding **z** and an amino acid sequence **a**. The encoding is processed by a MLP to yield a node embedding $\mathbf{h} \in \mathbb{R}^{L\times 256}$:

$$\mathbf{h} = \text{Linear}\big(\text{ReLU}\big(\text{Linear}(\mathbf{z}, in = 32, out = 256)\big), in = 256, out = 256\big).$$

The amino acid sequence is processed by an embedding layer to obtain an embedding $\mathbf{s}_{\text{aa}} \in \mathbb{R}^{L\times 32}$. Positional embeddings $\mathbf{p}_{\text{pos}} \in \mathbb{R}^{L\times L\times 128}$ are also created. An edge embedding $\mathbf{p} \in \mathbb{R}^{L\times L\times 128}$ is obtained by combining these tensors with:

$$\mathbf{p} = \text{LayerNorm}\Big(\big(\text{Linear}(\mathbf{s}_{\text{aa}}, in = 32, out = 128) \oplus \text{Linear}(\mathbf{s}_{\text{aa}}, in = 32, out = 128)\big) + \mathbf{p}_{\text{pos}}\Big),$$

where $\oplus$ denotes outer sum. We then provide **h** and **p** to a Structure Module as its single and pair representations, respectively, along with the amino acid sequence **a**. We adopt the Structure Module implementation from OpenFold[9]. We use 5 blocks and no dropout, as we found it to hurt reconstruction performance. All remaining hyperparameters are the default of OpenFold. The output of the final block is a 3D protein structure $\hat{\mathbf{R}} \in \mathbb{R}^{N_\text{h}\times 3}$ and is directly used as the output of the decoder.

#### 4 - Noise prediction network

##### 4.1 - Overview

**Outline**. The noise prediction network $\boldsymbol{\epsilon}_\theta$ has an architecture largely based on the Diffusion Transformer (DiT)[8,10]. The idea is to a create a representation **h** of the noisy input encoding and to updated it within the transformer using information from input conditions, embedded as node and edge embeddings **s** and **p**, respectively (**Supplementary Fig. 21a**).

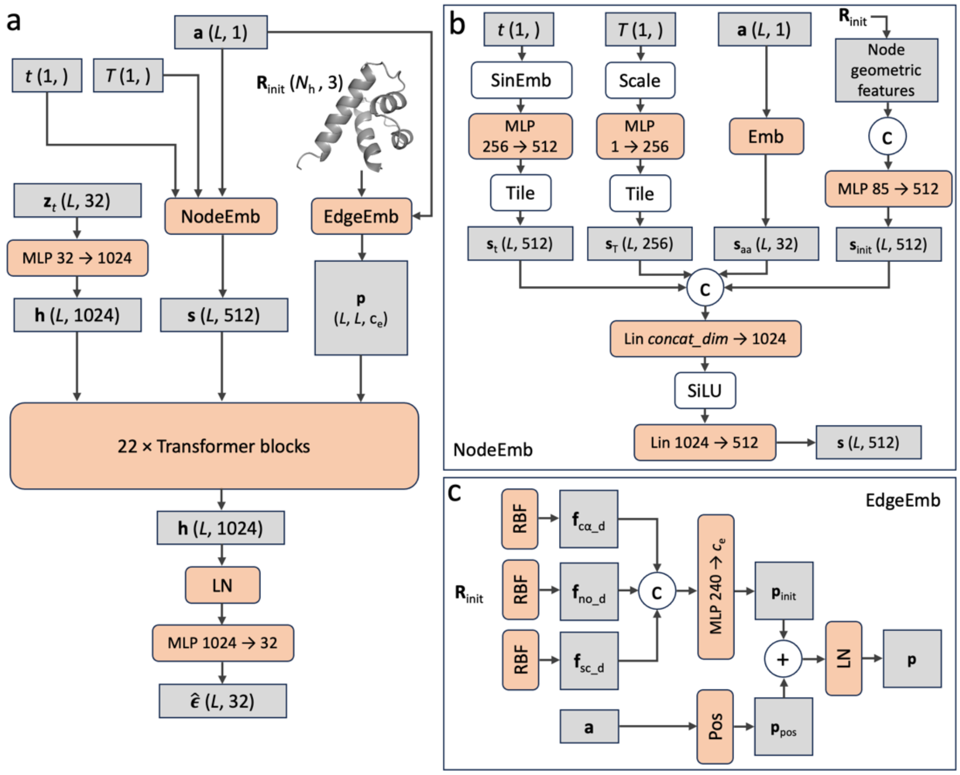


**Supplementary Fig. 20. Architecture of the noise prediction network of aSAM. a** Outline of the network. **b-c** Architecture of specific modules. **b** The temperature in is only used in the mdCATH-based model and so *concat_dim* in the second to last linear layer can vary depending on the model. SinEmb represent a sinusoidal embedding operation.

**Input**. The input of $\boldsymbol{\epsilon}_\theta$ is:

- $\tilde{\mathbf{z}}_t \in \mathbb{R}^{L\times 32}$: a noisy encoding at diffusion timestep $t$, representing a structure of a chain of length $L$.
- $t \in \mathbb{R}$: the timestep itself.
- $\mathbf{a} \in \mathbb{R}^{L\times 1}$: the amino acid sequence of the chain.
- $\mathbf{R}_{\text{init}} \in \mathbb{R}^{N_{\text{h}}\times 3}$: the heavy atom coordinates of the initial structure in the MD simulation of the chain.
- $T \in \mathbb{R}$: the simulation temperature. This input is used only in the aSAMt model trained on mdCATH.

#### 4.2 - Node representation

**Scaling the encodings**. To numerically help DDPM training, each of the 32 channels in an encoding $\mathbf{z} \in \mathbb{R}^{L\times 32}$ is standardized with the channel-wise mean and standard deviation in the dataset. The standardization happens prior to the addition of diffusion noise. At inference time, encodings generated by the DDPM are transformed back to their original scale, before being mapped to 3D structures by the decoder.

**Node representation**. A standardized noisy encoding $\tilde{\mathbf{z}}_t$ is processed via a MLP to yield a node representation $\mathbf{h} \in \mathbb{R}^{L\times 1024}$, which will be updated by a series of transformer layers.

#### 4.3 - Node conditional features

**From scalar inputs**. The timestep and temperature are converted into node features:

- $\mathbf{s}_{\mathrm{t}} \in \mathbb{R}^{L\times512}$: obtained by mapping $t$ on a sinusoidal embedding (with 256 dimensions), which is then processed with a MLP.

- $\mathbf{s}_{\mathrm{T}} \in \mathbb{R}^{L\times256}$: obtained by minmax scaling $T$ with $T_s = (T - T_{low})/(T_{up} - T_{low})$, then processing $T_s$ with a MLP. We set $T_{low} = 280\ K$ and $T_{up} = 470\ K$.

**From the input sequence**. The sequence **a** is embedded to $\mathbf{s}_{\mathrm{aa}} \in \mathbb{R}^{L\times32}$ via an embedding layer.

**From the initial structure**. $\mathbf{R}_{\mathrm{init}}$ is converted into node features that describe the local geometry of the chain:

- $\mathbf{f}_{\mathrm{c\alpha\_tor\_h}} \in \mathbb{R}^{L\times17}$: torsion angles formed by consecutive Cα atoms. The $L-3$ angles of a chain are bucketized into 16 equally sized bins from $-\pi$ to $\pi$, producing a one-hot encoding stored in the first channels. The last channel is filled with 0 for positions with missing angles and 1 for the rest.

- $\mathbf{f}_{\mathrm{c\alpha\_ba\_h}} \in \mathbb{R}^{L\times17}$: bond angles formed by consecutive Cα atoms. The $L-2$ angles of a chain are processed similarly to $\mathbf{f}_{\mathrm{c\alpha\_tor\_h}}$ into 17 channels.

- $\mathbf{f}_{\varphi\mathrm{\_h}} \in \mathbb{R}^{L\times17}$, $\mathbf{f}_{\psi\mathrm{\_h}} \in \mathbb{R}^{L\times17}$ and $\mathbf{f}_{\omega\mathrm{\_h}} \in \mathbb{R}^{L\times17}$: $\varphi$, $\psi$ and $\omega$ backbone torsion angles. For each angle type, the $L-1$ angles of a chain are processed similarly to the to $\mathbf{f}_{\mathrm{c\alpha\_tor\_h}}$.

These features are concatenated and mapped to a $\mathbf{s}_{\mathrm{init}} \in \mathbb{R}^{L\times512}$ tensor by an MLP.

**Node conditional embeddings**. $\mathbf{s}_{aa}$, $\mathbf{s}_{\mathrm{init}}$, $\mathbf{s}_{\mathrm{t}}$ and $\mathbf{s}_{\mathrm{T}}$ (if present) are concatenated and processed in the NodeEmb module to obtain $\mathbf{s} \in \mathbb{R}^{L\times512}$, the conditional node embeddings (**Supplementary Fig. 21b**).

##### 4.4 - Edge conditional features

**From the input sequence**. Positional embeddings $\mathbf{p}_{\mathrm{pos}} \in \mathbb{R}^{L\times L\times c_{\mathrm{e}}}$ are obtained from the sequence. In the ATLAS and mdCATH-based models $c_{\mathrm{e}}$ is set to 144 and 128, respectively.

**Edge features from the initial structure**. The coordinates $\mathbf{R}_{\mathrm{init}}$ are also converted into inter-atomic distances, used as edge features for the network:

- $\mathbf{f}_{\mathrm{c\alpha\_d}} \in \mathbb{R}^{L\times L\times144}$: an $\mathrm{RBF}(14.0, 144)$ of Cα-Cα distances.

- $\mathbf{f}_{\mathrm{no\_d}} \in \mathbb{R}^{L\times L\times32}$: an $\mathrm{RBF}(3.0, 32)$ of N-O distances (only in the ATLAS based-model).

- $\mathbf{f}_{\mathrm{sc\_d}} \in \mathbb{R}^{L\times L\times64}$: an $\mathrm{RBF}(7.0, 64)$ of distances between SCCs (only in the ATLAS-based model).

**Edge conditional embeddings**. All the available RBF expansions are processed together with $\mathbf{p}_{\mathrm{pos}}$ in the EdgeEmb module yield the $\mathbf{p} \in \mathbb{R}^{L\times L\times c_{\mathrm{e}}}$, the conditional edge embeddings (**Supplementary Fig. 21c**).

##### 4.5 - Transformer blocks

**Node representation updates**. The node representation **h** is updated through a series of 22 transformer blocks with the **s** and **p** embeddings providing conditional information. We use DiT blocks with an adaLN-Zero architecture[10] in which **s** is acting as a condition in the adaLN mechanism. We use 32 attention heads. To provide edge conditional information, at each block, **p** is linearly projected to attention biases used in the multihead self-attention update of **h**.

**Output**. After the last block, the **h** is converted to predicted noise $\hat{\boldsymbol{\epsilon}} \in \mathbb{R}^{L\times c}$ by an output module with an MLP.

#### 5 - aSAM training

##### 5.1 - Autoencoder loss

The per-sample loss $L_{\mathrm{AE}}$ used to train the AE is the sum of the following terms:

$$L_{\mathrm{AE}}(\mathbf{R}, \hat{\mathbf{R}}) = L_{\mathrm{rec}}(\mathbf{R}, \hat{\mathbf{R}}) + L_{\mathrm{nb}}(\hat{\mathbf{R}}) + L_{\mathrm{bl}}(\mathbf{R}, \hat{\mathbf{R}})$$

Where $\mathbf{R}$ and $\widehat{\mathbf{R}}$ are the original and reconstructed heavy atom positions of an MD conformation.

$L_{\text{rec}}$ a weighted sum of three terms directly adopted from the AF2 loss[7] (Algorithm 20 in the supplementary information in the original publication):

- FAPE loss over heavy atoms and all frames (line 28 in Algorithm 20), weighted by 0.75.
- FAPE loss over Cα atoms and backbone frames (line 17 in Algorithm 20), weighted by 0.75.
- Torsion angle loss (line 18 in Algorithm 20), weighted by 0.5.

Note that, like in AF2, the last two terms act both on $\widehat{\mathbf{R}}$, the final output of the Structure Module, and on intermediate reconstructions from every block of the module (which we all omit from the $L_{\text{AE}}$ definition). No other AE loss term of aSAM acts on intermediate structures.

$L_{\text{nb}}$ is defined as:

$$L_{\text{nb}}(\widehat{\mathbf{R}}) = \frac{1}{N_{\text{nb}}} \sum_{i,j} \max(m_{ij} - \hat{d}_{ij}, 0)$$

where the summation is over all pairs of residues $i$ and $j$ with $j - i \geq 3$, $N_{nb}$ if the number of such pairs, $\hat{d}_{ij}$ is the reconstructed distance between Cα atoms of $i$ and $j$ and $m_{ij}$ is a threshold distance specific to their amino acid types. The parameters of $m_{ij}$ are taken from the coarse-grained potential of the cg2all package[11].

$L_{\text{bl}}$ is defined as:

$$L_{\text{bl}}(\mathbf{R}, \widehat{\mathbf{R}}) = \frac{1}{N_{\text{bl}}} \sum_{i} |b_i - \hat{b}_i|$$

Where the summation is over all pairs of adjacent Cα atoms, $N_{bl}$ is the number of such pairs, $b_i$ and $\hat{b}_i$ are the "bonded" Cα-Cα lengths in the original and reconstructed 3D structures, respectively.

#### 5.2 - AE training protocol

When training the AE components we define two parameters: $n_{\text{systems}}$ and $n_{\text{frames}}$. The first is the total number of protein systems used in training, while the second is the number of snapshots uniformly sampled in a training epoch from MD data of each system (from all its trajectories). In other words, at each epoch the model sees a total of $n_{\text{systems}} \times n_{\text{frames}}$ samples. We train a model for a total of $n_{\text{epochs}}$. For the ATLAS dataset, each system corresponds to a different protein chain. For mdCATH, each system corresponds to a protein domain simulated at a certain temperature (therefore the same domain is found in 5 different systems). Note that in our setup, $n_{\text{systems}}$ can sometimes be lower than the total number of available systems in a dataset. In this case, at each epoch we randomly subsample $n_{\text{systems}}$ elements from the total amount. The ATLAS-based model is trained in two phases. In the first, we use only chains with length below 320 and set to zero the contribution of stereochemical loss terms. In the second, we increase the maximum length to 500 and add contribution from stereochemical terms. This helps minimize clashes and respect chain integrity, which FAPE alone did not fully resolve for larger proteins. For the mdCATH-based model, we ran only the first phase due to time constraints. We train with micro-batches containing proteins with the same number of residues to avoid padding strategies and we accumulate their gradients to use larger effective batch sizes. When training a model, we run multiple replicas and, at the end of training, we select the one with the best performance over the validation set. Depending on the training set, we use different number of replicas, $n_{\text{systems}}$, $n_{\text{frames}}$ , $n_{\text{epochs}}$ and batch size values. All models are trained with an Adam optimizer with $\beta_1 = 0.9$, $\beta_2 = 0.999$ and $\varepsilon = 1e - 8$. The starting learning rate is always 0.0005. We use a milestone schedule where the learning rate is multiplied by a factor of $\gamma = 0.5$ every $f_{\text{milestone}}$ epochs. Details of AE training processes and its computational times are summarized in **Supplementary Table 7**.

**Supplementary Table 7**. Training details of the AE models.

| | ATLAS phase 1 | ATLAS phase 2 | mdCATH |
|---|---|---|---|
| Protein length cutoff | $L \leq 320$ | $L \leq 500$ | $L \leq 320$ |
| Total training systems | 982 | 1174 | 25280 |
| $n_{\text{systems}}$ | 982 | 1174 | 17800 |
| $n_{\text{frames}}$ | 15 | 15 | 2 |
| $n_{\text{epochs}}$ | 240 | 240 | 160 |
| $w_{\text{nb}}$ | 0.0 | 0.01 | 0.0 |
| $w_{\text{bl}}$ | 0.0 | 1.0 | 0.0 |
| Initial learning rate (lr) | 0.0005 | 0.0005 | 0.0005 |
| Lr schedule | Milestone | Milestone | Milestone |
| Lr schedule parameters | $\gamma = 0.5$, $f_{\text{milestone}} = 40$ | $\gamma = 0.5$, $f_{\text{milestone}} = 40$ | $\gamma = 0.5$, $f_{\text{milestone}} = 20$ |
| Micro batch size | 2 | 1 | 2 |
| Batch size | 32 | 32 | 32 |
| GPUs | 2 × RTX2080 Ti | 2 × RTX2080 Ti | 2 × RTX2080 Ti |
| Total training steps | 110600 | 132200 | 178050 |
| Number of training replicas | 4 | 10 | 10 |
| Days for training a replica | 7 | 12 | 18 |

#### 5.3 – Denoising diffusion probabilistic model training protocol

When training the DDPMs we adopt a similar strategy and notations with respect to AEs (see above). Again, we use an Adam optimizer with $\beta_1 = 0.9$, $\beta_2 = 0.999$ and $\varepsilon = 1e - 8$. We adopt a "linear decay with warm up" learning rate schedule, which is commonly used for training transformer-like architectures. The learning rate starts at 0.0, linearly warms up to a baseline of 0.0005 in the first 10,000 steps, then linearly decreases for 615,000 steps until it reaches $5.0e - 6$. It finally stays as constant for the rest of training. We roughly stop training when the validation loss stops improving. Details of DDPM training processes are summarized in **Supplementary Table 8**. For information about the loss function and noise schedule of the DDPM, see the main text.

**Supplementary Table 8**. Training details of the DDPM.

| | ATLAS | mdCATH |
|---|---|---|
| Protein length cutoff | $L \leq 500$ | $L \leq 320$ |
| Total training systems | 1174 | 25245 |
| $n_{\text{systems}}$ | 1174 | 25245 |
| $n_{\text{frames}}$ | 100 | 5 |
| $n_{\text{epochs}}$ | 190 | 150 |
| Lr schedule | Linear with warm up | Linear with warm up |
| Base and minimum lr | 0.0005, 5.0e-6 | 0.0005, 5.0e-6 |
| Lr warmup steps | 10000 | 10000 |
| Lr decay steps | 615000 | 615000 |
| Micro batch size | 1 | 2 |
| Batch size | 32 | 32 |
| GPUs | 4 × RTX2080 Ti | 4 × RTX2080 Ti |
| Total training steps | 697800 | 593330 |
| Number of training replicas | 2 | 1 |
| Days for training a replica | 16 | 20 |

### Supplementary Note 2

#### Energy minimization in aSAM

aSAM frequently samples globally correct 3D structures, but with non-optimal side chain positions leading to clashes. For larger proteins or high temperatures (e.g.: $L > 300$ or $T > 400\,K$) there is also an increase in backbone atom clashes. To reduce clashes, we optimize the 3D structures generated by aSAM with an ad hoc energy minimization procedure implemented in PyTorch[12]. This framework allows optimization of multiple conformations in parallel, significantly improving computational efficiency.

#### 1 - Energy function

The energy function is based on: (i) terms from the Amber ff99SB force field[13], (ii) heuristic terms to minimize clashes and (iii) restraining terms derived from the snapshots generated by aSAM. The Amber ff99SB force field terms are reduced to a bare essential to obtain computational efficiency.

**Force field bond length potential.** For all covalent bond lengths in a protein, we use harmonic potentials $w_{bl}\left(b_i - b_i^{\mathrm{ff}}\right)^2$, where $b_i$ is a bond length in a conformation and $b_i^{\mathrm{ff}}$ is the energy minimum for that bond type from the force field. We do not use bond-specific force constants. The weight $w_{\mathrm{bl}}$ is set to $10^4$ for all bonds.

**Force field bond angle potential**. For all bond angles in a protein, we use harmonic potentials $w_{\mathrm{ba}}\left(\vartheta_i - \vartheta_i^{\mathrm{ff}}\right)^2$, where $\vartheta_i$ is a bond angle and $\vartheta_i^{\mathrm{ff}}$ is the force field minimum for that angle type. We do not use any angle-specific force constants. We set $w_{\mathrm{ba}}$ to $10^3$ for all angles.

**Force field proper torsion potential**. We only use selected Amber ff99SB proper dihedral terms for maintaining correct stereochemistry and planarity in the side chains in Arg, His, Phe, Trp and Tyr residues. We utilize a cosine potential $w_{\mathrm{pt}}\left(1 + \cos\left(n_i^{\mathrm{ff}}\alpha_i - \gamma_i^{\mathrm{ff}}\right)\right)$, where $\alpha_i$ is a torsion angle in a conformation and $n_i^{\mathrm{ff}}$ and $\gamma_i^{\mathrm{ff}}$ are the periodicity and phase parameters for that angle from the force field, respectively. Note that, since all angles that we consider have a single term in their cosine series expansion, we omit the summatory from the notation of the potential. We set $w_{\mathrm{pt}}$ to 10 for all angles.

**Force field improper torsion potential**. Similar to proper torsion angles, we only use those Amber ff99SB improper torsion terms for maintaining the correct planarity in the side chains of Arg, Asn, Asp, Gln, Glu, His, Phe, Trp and Tyr residues. Again, we employ a cosine potential $w_{\mathrm{it}}\left(1 + \cos\left(n_i^{\mathrm{ff}}\alpha_i - \gamma_i^{\mathrm{ff}}\right)\right)$, with all angles having a single term in their cosine series. The weight $w_{\mathrm{it}}$ is set to 10.

**Heuristic heavy atom repulsion potential**. To relax atomic clashes, we use an half-harmonic potential $w_{\mathrm{nb}}\max\left(d_{ij}^{\mathrm{lit}} - d_{ij}, 0\right)^2$, where $d_{ij}$ is a distance between two heavy atoms from separated by at least one position along the primary sequence and $d_{ij}^{\mathrm{lit}}$ is the literature sum of their van der Waals radii (with parameters from the *stereo_chemical_props.txt* file in the OpenFold distribution). The weight $w_{\mathrm{nb}}$ is set from 100 to 350 depending on the model (see below). To reduce memory consumption, we only apply the potential to a list of neighbor atoms. The neighbor list is created by selecting all heavy atoms from non-adjacent residues with a Cα-Cα distance below 10.0 Å. The list is first created from the input structure and then updated after every 10 minimization iterations.

**Restraints on adjacent Cα-Cα lengths**. We restrain all distances between pair of Cα atoms of adjacent residues along the primary sequence. We apply a harmonic potential $w_{\mathrm{ca}}\left(l_i - l_i^{\mathrm{ini}}\right)^2$ to any $l_i$ distance, with $l_i^{\mathrm{ini}}$ is the value of that distance in the structure directly generated by aSAM. We set $w_{\mathrm{ca}}$ to $10^5$. We adopt these terms, because the distribution of Cα-Cα lengths would otherwise lose its original width.

**Restraints on main chain torsion angles**. We restrain all $\phi$ and $\psi$ backbone torsion angles to the values observed in the initial aSAM snapshots. We apply a cosine potential $-w_{\mathrm{bb}}\left(1 + \cos\left(\alpha_i - \alpha_i^{\mathrm{ini}}\right)\right)$, where $\alpha_i$ is any $\phi$ and $\psi$ angle in a conformation and $\alpha_i^{\text{ini}}$ is its corresponding value in the structure directly generated by aSAM. The weight $w_{\text{bb}}$ is set to 50. Additionally, we apply a potential to $\omega$ angles. The term $w_{\text{om}}\left(1 + \cos(2\omega_i - \pi)\right)$ acts on all $\omega_i$ angles and forces them to be approximately flat. The weight $w_{\text{om}}$ is set to 10.

**Restraints on side chain torsion angles**. We restrain all $\chi_1$ to $\chi_4$ side chain torsion angles to the values in the SAM-generated snapshots. We apply a cosine potential $-w_{\text{sc}}\left(1 + \cos\left(\alpha_i - \alpha_i^{\text{ini}}\right)\right)$, where $\alpha_i$ is any $\chi$ angle in a conformation and $\alpha_i^{\text{ini}}$ is its value in the structure directly generated by aSAM. The weight $w_{\text{sc}}$ is set to $10^3$.

#### 2 - Optimization protocol

We initially developed an optimization protocol for the ATLAS-based aSAM and then updated it for the mdCATH-based model. This is because generating conformations at higher temperatures leads to more clashes and we needed a slightly more through optimization. For both protocols, we use a batch size of 50 conformations from the same protein.

**Optimization for the ATLAS-based model.** The protocol uses L-BFGS (limited-memory Broyden–Fletcher–Goldfarb–Shanno) optimization algorithm as implemented in PyTorch. We optimize the energy function described above, with the weight $w_{\text{nb}}$ for the heuristic atomic repulsion terms set to 100. We use a step size of 1.0 and run 20 optimization steps, with at maximum 10 iterations per optimization step.

**Optimization for the mdCATH-based model.** The protocol consists of a first stage using the Adam optimization algorithm as implemented in PyTorch, and a second stage using the L-BFGS algorithm. We optimize the energy function described above and set $w_{\text{nb}}$ to 100 and 250 in the first and second stages, respectively. In the first stage, we use a step size of 0.001, set the Adam $\beta_1$ and $\beta_2$ parameters to 0.5 and 0.9, and run 50 optimization steps. In the second stage, we use a step size of 1.0 and run 30 optimization steps, with at maximum 10 iterations per step. To maintain a balance between computational efficiency and the quality of optimization outcome, after every iteration we count the number of clashes in the neighbor list, defined as distances $d_{ij} \leq 1.75$ Å. If, at any point in the two stages, the average number of clashes in a batch surpasses a threshold $n_{\text{ct}} = 0.7$, we stop the energy minimization of that batch. For small proteins and low temperatures, optimization is typically stopped at the beginning of the first stage. For larger proteins and higher temperatures, optimization often runs through all of the two stages. When generating melting curves with aSAM, we set always set $n_{\text{ct}} = 0.3$ to help remove additional clashes that may occur at out-of-training temperatures.

### Supplementary Note 3

### Comparison with experimental data

#### 1 - Protein monomers with experimentally measured $T_m$

**Protein selection.** The 62 proteins used in **Fig. 7a** were selected from the $S^{[Temp]}$ dataset by Pucci et al.[14], which contains 222 proteins with an experimentally measured $T_m$ and a structure on the PDB. aSAMt was trained on mdCATH simulations, which contain monomeric proteins with no disulfide bridges or ligands. To obtain a meaningful comparison between the model predictions and experimental data, we removed from the dataset all proteins meeting any of the following criteria:

- The biological unit reported in the dataset is not a “monomer”. Oligomeric contacts play a crucial role in protein stability. We also excluded the following entries: 1J2V since it is described as Homo 3-mer on the PDB, and 4TLN since the publication reporting the $T_m$ measurement describes the domain as a stable dimer.
- Membrane proteins (e.g: 1COL).
- Proteins with the following stabilizing structures or ligands: any disulfide bridges; proteins bound to heme ligands (e.g.: myoglobins or cytochromes); proteins stabilized by iron-sulfur clusters (e.g.: ferrodoxins) or coordinating calcium ions (e.g: calmodulins).
- Any protein for which the $T_m$ measurement was performed at pH < 4.5. Low pH impacts protein stability.
- Proteins with more than 700 residues, since the maximum length of training proteins for aSAMt is 320. For such large proteins, ensembles generated at > 450 K often contain too many clashes.

Additionally, we removed all proteins with more than 45% sequence identity with any mdCATH training domain. After applying all these filters, we obtained a list of 62 proteins with the following PDB codes:

1PIN-Nterm, 1SHG, 1C8C, 1AZP, 1CSP, 1MJC, 2HPR, 1HA4, 1Y4Y, 2BJD, 1TEN, 1FNA, 1IV7, 1AG6, 1H7M, 2OV0, 1EW4, 1TMY, 1FGA, 2TDX, 5DFR, 1BD8, 1ZDR, 2LZM, 1AMM, 1F0L-Nerm, 1JNX, 1P3J, 1IO2, 1BMC, 1ANK, 1S3G, 1ZIP, 1WDN, 4U2B, 1CHK, 1MVE, 1I4N, 4BLM, 1WQ5, 1SVN, 1URP, 2ST1, 1SUP, 1MJ5, 1ABE, 1EVQ, 1JJI, 1AVR, 1CEC, 2CNC, 1KE4, 2POO, 1GTG, 1IS9, 3MBP, 1ATN, 1QLP, 1EFC, 3PGK (1QPG), 1PII, 1G5A

**Input structures for aSAMt.** Whenever possible, we used as input to aSAMt raw PDB structures. For entries with missing atoms or residues, we fixed their PDB structures with MODELLER[15]. We substituted 3PGK with 1QPG, a more recent entry of the same protein. For 1PIN-Nterm, 1C8C, 1AZP and 1CHK we used an AlphaFold2 model as input. 1PIN-Nterm is an N-terminal fragment of a larger protein interacting with other domains, 1C8C and 1AZP are small proteins bound to a DNA fragment, and 1CHK was determined at pH 5.5 but its $T_m$ was measured at pH 7.0. The model was trained with MD snapshots of monomers as input condition (**Methods**) and we decided to use AF2 models since we expected them to be more similar to training input with respect to such raw PDB structures.

#### 2 - Homologous proteins with experimentally measured $T_m$

**Protein selection.** The 5 pair of homologous proteins described in **Supplementary Table 6** were obtained from the dataset in Razvi and Schotlz[5]. To select the proteins, we adopted the seven criteria listed above.

**Input structures for aSAMt.** Whenever possible, we used as input to aSAMt raw PDB structures. Structures with missing atoms or residues, were fixed with MODELLER. We substituted 3PGK with 1QPG, a more recent entry. Similar to what we described above, for 1G6P (an NMR structure), 1RIL (a 2.8 Å resolution X-ray structure), and 1CN7 (an NMR structure) we used AF2 models as input to aSAMt. To use similar input conditions, we also used an AF2 models for their homologous protein.
